# Molecular evidence of chemical disguise by the socially parasitic spiny ant *Polyrhachis lamellidens* (Hymenoptera: Formicidae) when invading a host colony

**DOI:** 10.1101/2022.05.22.492802

**Authors:** Hironori Iwai, Masaru Mori, Masaru Tomita, Nobuaki Kono, Kazuharu Arakawa

## Abstract

While most ant species establish a colony independently, some socially parasitic ants build the foundation of their colony by invading other ant (host) colonies and utilizing their labour forces. Many socially parasitic ants disguise their cuticular hydrocarbon (CHC) profile, which is also known as nestmate discrimination pheromones, when invading the host colony. Since the strategy of chemical disguise is widespread in socially parasitic ants, elucidating the mechanism of chemical disguise will promote knowledge on the evolutionary history of social parasitism. However, detailed knowledge is still lacking, as the relevant information has only originated from circumstantial evidence, which was obtained from ecological observations. In this study, we investigated the mechanism of chemical disguise in a new queen of a temporary socially parasitic spiny ant (*Polyrhachis lamellidens*) by measuring its CHC profile, performing a tracing assay with labelled substances, and analysing gene expression levels. First, after rubbing behaviour was observed against the host workers, the CHC profile in *P. lamellidens* shifted to pronounced peaks that closely resembling that of the host workers. We also observed a reduction in aggressive behaviours by the host ant against *P. lamellidens* after rubbing behaviour was performed. In addition, *P. lamellidens* acquired artificially-applied labelling substances from host workers through their rubbing behaviours, while gene expression profiling showed the expression of CHC synthesis-related genes did not change during this behaviour. These results suggest that *P. lamellidens* directly obtains host CHCs through rubbing behaviour, and these host CHCs enables *P. lamellidens* to remain disguised during colony invasion.

## 1 Introduction

There are several hydrocarbons present on ant cuticles, and these hydrocarbons function as semiochemicals (Detrain et al., 1999; Howard & Blomquist, 2005; Sturgis & Gordon, 2012). The cuticular hydrocarbon (CHC) profile of ants varies among different colonies. Ants discriminate between nestmates and foreign enemies (non-nestmates) by recognizing the differences in CHC profiles and behave aggressively towards individuals with different CHC profiles, even if the non-nestmates are the same species. On the other hand, organisms that evade the nestmate discrimination by ants and utilize the work force of ants are collectively referred to as “myrmecophiles” or “social parasites” (Buschinger, 2009; Kistner, 1982). It is known that some myrmecophile species disguise their own CHC profile to obtain a similar profile to that of the host. It is believed that myrmecophiles utilize chemical disguises to avoid being exclusively discriminated as non-nestmates by host ants (Akino, 2008; Dettner & Liepert, 1994; Lenoir et al., 2001). For example, when a newly mated queen in the slave-making parasitic ant Polyergus (Hymenoptera: Formicidae) enters a host colony, she temporarily avoids aggressive behaviours from the host worker by emitting a repellent substance, then she kills the host queen, disguises her CHC profile, and exhibits the CHC profile of the host queen (d’Ettorre & Errard, 1998; d’Ettorre et al., 2000; Johnson et al., 2001; Mori et al., 1995; Tsuneoka, 2008; Tsuneoka & Akino, 2009 & 2012; Zimmerli & Topoff, 1994).

There are two methods of chemical disguise, including the chemical camouflage and chemical mimicry methods (Akino, 2008; Dettner & Liepert, 1994; Lenoir et al., 2001). The former method is utilized by organisms to acquire the CHC profile from the host directly, and the latter is utilized to biosynthesize host-like CHC. Some studies have experimentally verified the mechanism of chemical disguise in several myrmecophile species, such as spiders and crickets (Akino et al., 1996; Howard et al., 1990; Scarparo et al., 2019; von Beeren et al., 2011 & 2012; Vander Meer & Wojcik, 1982). The mechanism of chemical disguise was estimated in socially parasitic ants through ecological observations (Bauer et al., 2010; Lenoir et al., 1997). However, there is little information on the quantitative and molecular mechanisms that describe the intricacies of chemical disguise. Since the strategy of chemical disguise is a feature that is unique to socially parasitic ants, we believe that elucidating the molecular basis of the detailed mechanisms will elucidate the evolutionary history of social parasitism.

*Polyrhachis lamellidens* is an appropriate species to utilize when investigating the detailed mechanism of chemical disguise. *P. lamellidens* is an ant species that is temporary socially parasitic and utilizes several species of hosts, including *Camponotus japonicus* and *Camponotus obscuripes*, and the presumed candidate host, *Camponotus kiusiuensis* (Iwai et al., 2021; Japanese Ant Database Group, 2003; Kohriba, 1963 & 1966; Kubota, 1974; Kurihara et al., 2022; Sakai, 1990, 1996, & 2000; Yano, 1911). In this species, the newly mated queen performs distinct behaviours when invading the host colony, and these behaviours include straddling the host worker and rubbing its body with her legs (Japanese Ant Database Group, 2003; Kohriba, 1963; Kubota, 1974; Kurihara et al., 2022; Sakai, 1990 & 2000) (Figure 1A). This behaviour, which is known as a “rubbing behaviour”, is unique to *P. lamellidens* queen that are newly mated, is considered a strategy of chemical disguise, and prevents attacks by the host (Kohriba, 1963); however, little is known about the molecular machinery of this strategy.

In this study, we quantitatively evaluated the efficiency of rubbing behaviour-induced chemical disguise by measuring CHC profiles and conducting behavioural tests, and this was performed to clarify the effect of rubbing behaviour on the CHC of *P. lamellidens* and nestmate recognition of host workers. Furthermore, we elucidated the mechanisms of the disguise strategy of *P. lamellidens* by performing tracing assays with the labelled substances and by analysing the expression of related genes, aiming to reveal the behavioural ecology, biochemistry, and molecular biology foundations that explain social parasitism in ants. Therefore, in this study, GC-MS analyses of CHC were performed to quantitatively evaluate the extent to which the rubbing behaviour achieves chemical disguise. Through behavioural experiments, we examined the reactions of host workers to the chemical disguise obtained by rubbing behaviour. Furthermore, by labelling assays and gene expression profiling, we investigated whether chemical disguise is implemented by camouflage or mimicry.

**Figure 1.**
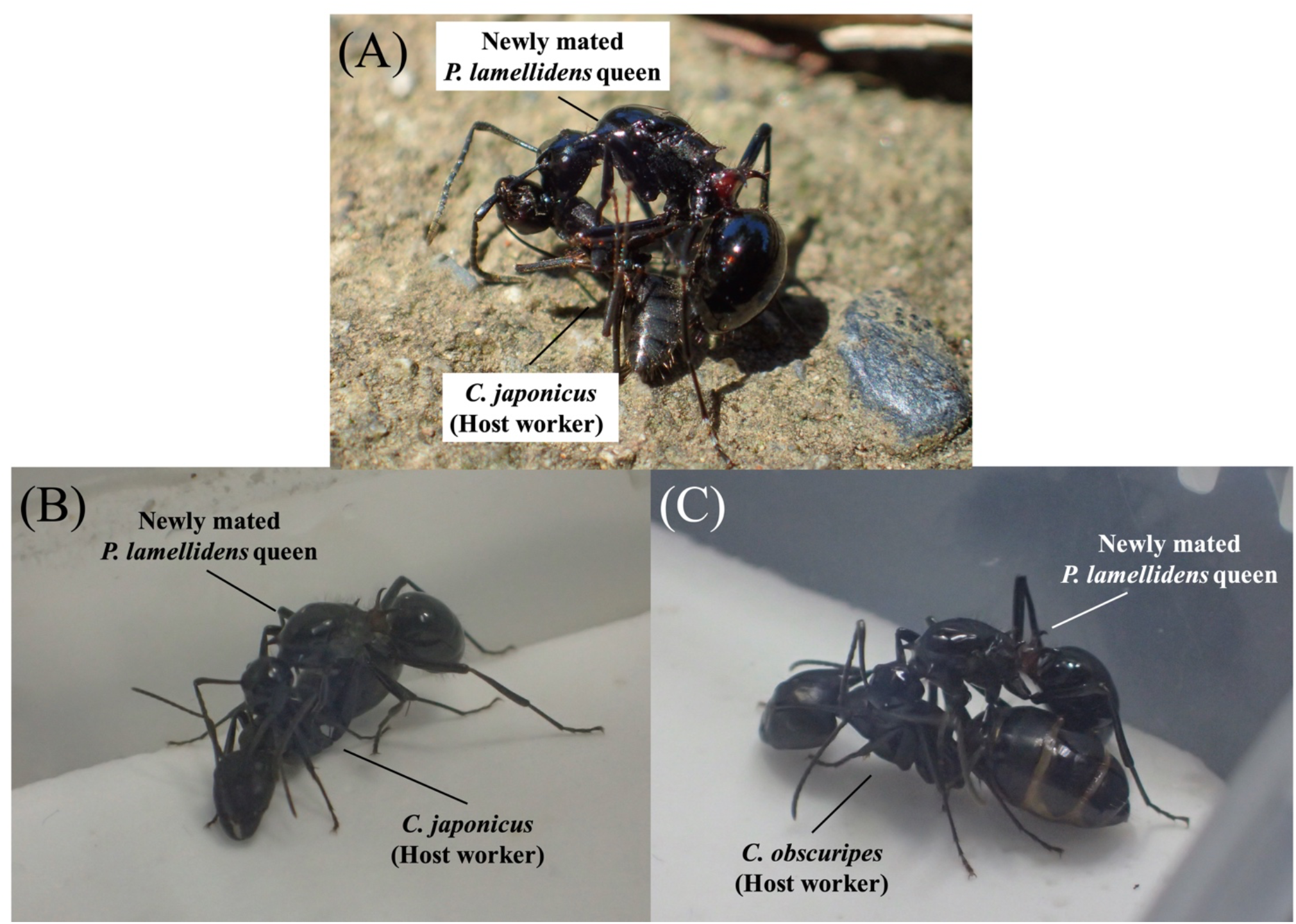
The rubbing behaviour of newly mated *P. lamellidens* queens. (A) The newly mated *P. lamellidens* queen exhibits rubbing behaviour towards its host worker (*C. japonicus*) in a natural environment at a mixed forest in Nagaoka City, Niigata Prefecture (N 37°25′51′′, E 138°52′53′′). (B) Rubbing behaviour against *C. japonicus* workers under laboratory conditions. (C) Rubbing behaviour against *C. obscuripes* workers under laboratory conditions.

## 2 Materials and Methods

### 2.1 Sampling and rearing

Ant samples were collected in Niigata, Yamanashi, and Yamagata Prefecture, Japan (2015 to 2021). Approximately 50 newly mated *P. lamellidens* queens were collected from a mixed forest in Nagaoka City, Niigata Prefecture (N 37°25′51′′, E 138°52′53′′), and after the queens finished their nuptial flights, their original colony was not known. A *P. lamellidens* colony was collected from a primeval beech forest in Nirasaki City, Yamanashi Prefecture (N 35°42′49′′, E 138°29′12′′); the colony approximately consisted of three queens, more than 1,000 workers and larvae. *C. japonicus* and *C. obscuripes* colonies were collected from a primeval beech forest in Nagaoka City, Niigata Prefecture (N 37°41′42′′, E 138°88′95′′), a mixed forest in Tsuruoka City, Yamagata Prefecture (N 38°33′49′′, E 139°55′31′′), and a mixed forest in Minamikoma District, Yamanashi Prefecture (N 35°46′11′′, E 138°29′43′′); the average colonies consisted of 100 workers for *C. japonicus* and one queen and 200 workers for *C. obscuripes*. The host workers used in each experiment were from the same colony each time.

The rearing method was adjusted to the colony size. Newly mated *P. lamellidens* queens were reared individually in a plastic tub that was 4.5 cm in length, 2.5 cm in width, and 2.0 cm in height and contained moistened tissue. Each collected ant colony was reared separately in a plastic tub (17.5 cm in length, 8.0 cm in width, 3.0 cm in height, or 20.5 cm in length, 10.5 cm in width, 7.0 cm in height) in which plaster was laid and a feeding area was established. Newly mated *P. lamellidens* queens were fed 5 μl of 50% maple syrup (Maple Farms Japan, Osaka, Japan) every 7–10 days. Ant colonies were also fed 100–5000 μl of 50% maple syrup every 7–10 days. Worker ants were also fed mealworms, crickets, cockroaches, and/or 8–16 g of Pro Jelly for beetles and stag beetles (Wraios, Saitama, Japan). Newly mated *P. lamellidens* queens were reared in a cool incubator (Mitsubishi Electric Engineering Company, Tokyo, Japan) under dark (0 L:24 D) conditions at 15–20°C. *C. japonicus* and *C. obscuripes* were reared under light-dark conditions (14 L10 D) at 20–30°C. Different rearing conditions were adopted for each species due to our previous rearing experience. In particular, newly mated *P. lamellidens* queens were observed to be sensitive to high temperatures, which is why we adopted the above rearing method.

### 2.2 Induction of rubbing behaviour under laboratory conditions

The newly mated *P. lamellidens* queen and host workers (*C. japonicus* or *C. obscuripes*) were placed in plastic cases that contained plaster (5.0 cm x 4.5 cm x 2.5 cm) (Figure 1, Supplementary Movie 1). To prevent possible attacks by the host worker before the newly mated *P. lamellidens* queen performed rubbing behaviours, the host worker was anesthetized through freezing (it was exposed to an environment of 4°C for 2 min and -20°C for 3 min) before the experiment. A newly mated *P. lamellidens* queen could perform the rubbing behaviour for three days. When the newly mated *P. lamellidens* queens continued the rubbing behaviour for more than one day, we added one new host worker (all from the same colony) every day until the third day. Host workers that spent one week in the nest were used this experiment. The newly mated *P. lamellidens* queens that were prepared in this process were used for the various experiments that followed. The individuals that performed rubbing behaviour at least once were used for later experiments. In the isolation rearing, only the host workers were removed from the case, and they were maintained for one to nine days after the rubbing behaviour was observed. We conducted the rubbing behaviour and isolation rearing experiments under light conditions (24 L:0 D) at 20–30°C and did not feed the ants throughout these experiments. As a control, we also prepared a newly mated *P. lamellidens* queen without exposure to host workers.

### 2.3 Observations of host worker aggressions against the newly mated *P. lamellidens* queen by behavioural tests

To observe the aggressive behaviours of host workers on the newly mated *P. lamellidens* queen, all host workers were removed immediately after rubbing behaviour had occurred for three days, and the newly mated *P. lamellidens* queen was transferred to a plastic cup that was lined with plaster (7.6 cm in diameter and 3.8 cm in height). Three host workers from the same colony as that used in the rubbing behaviour experiment were transferred into a plastic cup that contained a newly mated *P. lamellidens* queen that already exhibited rubbing. The first 10 behaviours that host workers performed within 1 min after being transferred to the plastic cup were recorded according to the following definitions:

- Ignore: the ant does not respond aggressively upon contact with the target.
- Threaten: the ant opens its mandibles or maintains distance from the target.
- Rush: the ant approaches and quickly bites the target.
- Bite: the ant continuously bites the target for at least one second.

Among these behaviours, “threaten”, “rush”, and “bite” were considered aggressive behaviours. The host workers were selected at random and were used only once in each test. The host workers used for the test were transferred to another case and were not used in subsequent tests.

### 2.4 Application of labelled substances for the tracing assay

We used the stable isotopes of n-triacontane (C_30_H_62_) and n-dotriacontane (C_32_H_66_) as labelled substances, and these substances have similar carbon chain lengths to those of the ant CHC (CAS: 93952-07-9, 62369-68-0). Each substance was dissolved individually in hexane (16 mg/mL), 100 µl of the solution was added to a 300 µl micro insert (GLC4010-S630; Shimadzu, Kyoto, Japan), and the solvents were evaporated with nitrogen gas. The solution was stirred with an end-to-end pipette, during which nitrogen gas was sprayed to evenly apply the labelled substances over the entire inner wall of the insert. After the solvent was completely volatilized, the labelled substances were applied to the host body surfaces without inducing injury, and this was achieved by placing the host worker into the insert where the labelled substances (C_30_D_62 &_ C_32_D_66_) were applied to the inner wall and gently shaking the hosts for 1 h using a rotator.

### 2.5 Extraction and analysis of hydrocarbons

The method for extracting cuticular hydrocarbons was modified from a previous study (Akino & Yamaoka, 2012). A single individual was exposed to 4°C for 2 min and then -20°C for 3 min for anaesthetic purposes and was then immediately placed in a disposable glass tube (trunk diameter: 10 mm, total height: 75 mm) that contained 200 μl of hexane mixed with saturated alkane (docosane: C_22_H_46_; 10 ng/µl) as an internal standard material. The chemical materials from the body surfaces of ants were extracted by dipping the ants in hexane for 5 min. After separation in 1X PBS, the chemical materials from the postpharyngeal glands were extracted by being dipped in hexane and were sonicating using a Bioruptor II (BM Equipment, Tokyo, Japan). Next, 0.5 g of silica gel C-200 (Wako Pure Chemical Industries, Tokyo, Japan) was used to fractionate hydrocarbons from the extracted materials. After washing with hexane, silica gel was placed inside a glass pipette that was installed on the stand. Fractionation was carried out by adding the extract inside the pipette. At this time, 1 ml of hexane was added into the pipette to elute hydrocarbons. The obtained hydrocarbons were concentrated by applying nitrogen gas until the solvent hexane was completely evaporated. Thereafter, 50 μl of hexane was added to the vial, and the concentrated cuticular hydrocarbon was eluted.

The GC-MS system comprised an Agilent 6890N, and an Agilent 5973 MSD was used to measure hydrocarbons. The column used for MS was an HP-5 MS (length 30 m, diameter 0.250 mm, film thickness 0.25 μm; Agilent Technologies, California, USA). Two microlitres of sample was injected. The splitless mode was adopted for the sample injection port and the apparatus was maintained at 300°C. Helium was used as the carrier gas at a constant flow rate setting of 0.9 ml/min. The oven temperature was set as follows: 40°C for 3 min, 40 to 260°C at 30°C/min, 260 to 300°C at 15°C/min, and 18 min at the final temperature. C_7_ to C_40_ saturated alkanes were used as standard substances.

The estimation of cuticular hydrocarbons and statistical analysis were carried out based on peak areas. A previous study (Ozaki et al., 2005) was referred to estimate the CHC of *C. japonicus*. The peak areas were transformed according to Z_ij_=ln[Y_ij_/g(Y_j_)], where Y_ij_ is the peak area _i_ for the individual _j_ values and g(Y_j_) is the geometric mean of all peak areas for the individual _j_ values (Reyment, 1989). Hierarchical clustering analysis and correlation analysis were performed using a standardized data matrix of CHC to investigate the similarity between parasite and host CHC profiles. In the hierarchical clustering analysis, the Euclidean distance between each sample was calculated and clustered by the Ward method. In the correlation analysis, the Pearson correlation coefficient was calculated, and the similarity of the CHC profile between each group was evaluated. Statistical analysis was carried out using R software v 3.2.0 (R Core Team, 2015; http://www.R-project.org/), and the gplots package was used to create a heatmap (Warnes et al., 2016; http://CRAN.R-project.org/package=gplots).

### 2.6 Total RNA extraction

Total RNA extraction was carried out for the whole body (5 larvae, 1 worker, and 1 queen), abdomen (18 newly mated queens), and fat body in the abdomen (17 newly mated queens) of *P. lamellidens* according to a previous study (Kono et al., 2016). Samples were exposed to liquid nitrogen and were placed in a ZR BashingBead lysis tube (Zymo Research, California, USA) that contained 600 µl of TRIzol Reagent (Life Technologies, California, USA). The samples were crushed with a Multi-Beads Shocker at 2,500 rpm for 30 sec (Yasui Kikai, Osaka, Japan), and total RNA was extracted with a Direct-zol RNA MiniPrep (Zymo Research, California, USA) without DNase treatment. The extraction of total RNA from the abdomen fat body was performed according to Koto et al., 2019. The abdomen fat body was dissected from the newly mated *P. lamellidens* queens in ant saline [4.8 mM TES, which contained 127 mM NaCl, 6.7 mM KCl, 2 mM CaCl2, and 3.5 mM sucrose] and placed in BioMasher II (Nippi, Tokyo, Japan) that contained 110 µl TRIzol Reagent. After the tissue was disrupted in BioMasher II, extraction was performed with a DNase treatment using a Direct-zol RNA MicroPrep (Zymo Research, California, USA). We performed a quality check (RIN, quantity, purity ratio) for the total RNA by utilizing TapeStation 2200 RNA Screen Tape (Agilent Technologies, California, USA), Qubit Broad Range (BR or HS) RNA assay (Life Technologies, California, USA), and NanoDrop 2000 (Thermo Fisher Scientific, Massachusetts, USA).

### 2.7 Library preparation and cDNA sequencing

Illumina sequence libraries were prepared with the extracted total RNA using the NEBNext Ultra RNA Library Prep Kit for Illumina (New England BioLabs, Massachusetts, USA) or KAPA mRNA HyperPrep Kit (KAPA BIOSYSTEMS, North Carolina, USA). Libraries that contained the whole body and abdomen were prepared using the NEBNext Ultra RNA Library Prep Kit for Illumina. The mRNAs of the whole body and abdomen were isolated by NEBNext Oligo d(T)25 beads (New England BioLabs, Massachusetts, USA) from 100–200 ng of total RNA. ds cDNA was synthesized from the isolated mRNA using ProtoScript II Reverse Transcriptase and NEBNext Second Strand Synthesis Enzyme Mix (New England BioLabs, Massachusetts, USA). The synthesized cDNAs were end-repaired by NEBNext End Prep Enzyme Mix and added to NEBNext Adapter for Illumina (New England BioLabs, Massachusetts, USA). The addition adapter to cDNA and amplification were achieved through PCR (16 cycles).

The mRNAs of ant fat bodies were isolated by mRNA Capture Beads (KAPA BIOSYSTEMS, North Carolina, USA) from 100–200 ng of total RNA. ds cDNA was synthesized from the isolated mRNA using KAPA Script and 2^nd^ Strand Synthesis & A-Tailing Enzyme Mix (KAPA BIOSYSTEMS, North Carolina, USA). The synthesized cDNAs were end-repaired by 2^nd^ Strand Synthesis & A-Tailing Enzyme Mix and Adapter Ligation Master Mix (KAPA BIOSYSTEMS, North Carolina, USA). The adapter addition to cDNA and amplification were achieved through PCR (16–18 cycles).

The cDNA libraries were sequenced by the NextSeq 500 (Illumina, California, USA) in pairs or single-ends with 150 or 75 cycles of the NextSeq 500/550 High Output Kit v2.0 (Illumina, California, USA). We evaluated the accuracy of sequence reads by FastQC v0.11.9 (http://www.bioinformatics.babraham.ac.uk/projects/fastqc/).

### 2.8 De novo transcriptome assembly and gene prediction

The de novo assembly of the transcriptome sequence reads was performed with Trinity v2.8.5 (Grabherr et al., 2011). We performed a quality check of the constructed transcriptome assembly by evaluating BUSCO v2/v3 (reference gene set: Arthropoda) using gVolante v1.2.1 (Nishimura et al., 2017). We used Augustus v3.2.2 to predict the gene regions that were present on the transcriptome assembly using the *Camponotus floridanus* gene model, a closely related species (Stanke & Morgenstern, 2005). CD-HIT-EST (Huang et al., 2010) removed the predicted gene regions with overlapping sequences in the cut-off value of 0.9.

### 2.9 Gene annotation and expression analysis

We annotated CHC synthesis-related genes in the transcriptome assembly of *P. lamellidens* with predicted coding regions. The CHC synthesis-related genes (fatty acid synthase, desaturase, elongase, cytochrome P450 decarbonylase) were annotated using the *Drosophila melanogaster* genes that were registered in UniProt as a query (Q9VQL6_DROME, M9PB21_DROME, ELOF_DROME, Q7K4Y0_DROME, Q9VG68_DROME, A7DZ97_DROME, CP4G1_DROME). After searching for candidate genes in the transcriptome assembly of *P. lamellidens* using BLAST similarity search (E-value < 1E-30; Camacho et al., 2009), the obtained candidate genes and query sequences were subjected to a protein domain search using HMMER v3.1b2 (Eddy, 2011) with the Pfam-A database (E-value ≤ 1E-10). The domain structure was confirmed using DoMosaics v0.95 (Moore et al., 2014). Gene expression was quantified as transcripts per million (TPM) using Kallisto v0.43.0 (Bray et al., 2016). The genes that possessed a domain structure similar to that of *D. melanogaster* and exhibited a TPM value of 10 or higher in any sample were considered to be CHC synthesis-related genes of *P. lamellidens*. We used the R package Edge R v3.18.1 with an FDR < 5% for searching differentially expressed genes (DEGs) (Robinson et al., 2010). We also annotated DEGs by a tBLASTn similarity search using the nr database.

### 2.10 RT–qPCR

We synthesized cDNA from 100 ng of total RNA extracted from the fat body in the abdomens using SuperScript III Reverse Transcriptase (Invitrogen, Massachusetts, USA). qPCR was performed using a KAPA SYBR Fast qPCR Kit (KAPA BIOSYSTEMS, North Carolina, USA) and LightCycler 96 (Roche, Basel, Switzerland). The primer list is provided in Supplementary Table 1. Preliminary tests confirmed that the amplification efficiency of all primers was similar (1.8∼2.2). We used the housekeeping genes *Gapdh1* and *Actin-5C* as reference genes. The amplicon lengths of all genes were between 143–163 bp. We calculated the relative expression level of target genes by the E-Method using LightCycler 96 Application software (Roche, Basel, Switzerland).

### 2.11 Protein expression analysis

The fat body from a newly mated queen of *P. lamellidens* was dissected in ant saline. The tissue was crushed in lysis buffer (12 mM sodium deoxycholate, 12 mM sodium N-dodecanoylsarcosinate, and 50 mM ammonium bicarbonate containing 1% protease inhibitor cocktail for general use [Nacalai Tesque, Kyoto, Japan]) followed by sonication for 20 min. The lysate (20 µg protein/50 µl) was reacted with 0.5 µl of 1 M dithiothreitol at 37°C for 30 min followed by 2.5 µl of 1 M iodoacetamide at 37°C for 30 min in the dark. After adding 200 µl of 50 mM ammonium bicarbonate, the sample was digested using 0.6 µg of Lys-C (Wako Pure Chemical Industries, Tokyo, Japan) at 37°C for 3 h followed and 0.5 µg of trypsin (Promega, Madison, WI, USA) at 37°C for 16 h. The digest was acidified with trifluoroacetic acid and sonicated for 10 min. The supernatant was desalted using C18-StageTips (Rappsilber et al., 2003) and dried under reduced pressure.

To perform proteome analysis, a system equipped with a nanoElute and a timsTOFPro (Bruker Daltonics, Bremen, Germany) was used. The digests that were dissolved in 0.1% formic acid 2% acetonitorile (0.4 µg/µl) were injected into a spray needle column (ACQUITY UPLC BEH C18, 1.7 µm, ID = 75 µm, length = 25 cm) and were separated by gradient analysis. Mixtures of (A) formic acid/water (0.1/100, v/v) and (B) formic acid/acetonitorile (0.1/100, v/v) were used as the mobile phase. The composition of the mobile phase (B) was changed to 2%–35% in 100 min and 35%–80% in 10 min while maintaining a flow rate of 280 nl/min at 60 °C.

Tandem MS was performed using a parallel accumulation serial fragmentation (PASEF) scan mode (Meier et al., 2018). Briefly, the peptides were ionized to positive ion at 1,600 V. Trapped ion mobility spectrometry scanning was performed at a 1/K0 range of 0.7–1.2 Vs/cm2 with a ramp time of 100 ms while maintaining the duty cycle at 100%. MS scanning was performed at a mass range of m/z 300– 1,200 followed by 10 PASEF-tandem MS scans per cycle (precursor ion charge = 0–5, intensity threshold = 500, target intensity = 2,000, isolation width = 2 Th at m/z 700 and 3 Th at m/z 800, collision energy = 20 eV at 0.6 Vs/cm2, 59 eV at 1.6 Vs/cm2). The raw data and the associated files were then deposited in the ProteomeXchange Consortium via the jPOST partner repository (accession number: JPST001552) (Okuda et al., 2017).

Protein identification and quantitative analysis of CHC synthesis-related proteins were performed using FragPipe v15.0 containing MSFragger v3.2 and Philosopher v3.4.13 (Kong et al., 2017). The protein sequence database was generated from transcriptome assembly, and the decoy reversed sequences were added. We also used the cRAP database (https://www.thegpm.org/crap/) to detect contaminant proteins.

The mass tolerance of the precursor and the fragment ions was set to 20 ppm and 0.05 Da, respectively. Mass calibration and parameter optimization were performed using the Philosopher algorithm (da Veiga Leprevost et al., 2020). The enzyme was set to trypsin as a specific cleavage, and up to two missed cleavages were allowed in the proteolysis. The allowed peptide lengths and mass ranges were 7–50 residues and 500–5,000 Da, respectively. Carbamidomethylation was set as a fixed modification at the cysteine residue, whereas N-acetylation at the protein N-term and oxidation at the methionine residue were set as variable modifications, and this allowed for up to three sites per peptide. The peptide spectrum matches and the identified peptides/proteins were determined at <1% FDR at the protein level.

Quantitative analysis of each peptide was performed with matches between runs (m/z tolerance = 20 ppm, retention time tolerance = 3 min, ion mobility tolerance = 0.05 Vs/cm2). After removing outlier values for the identified peptides, the intensities of the identified peptide in the data were normalized (the median log2 values were unified). The MaxLFQ algorithm (Cox et al., 2014) was used to compare the levels of protein expression between runs using the normalized intensity values of identified unique peptides.

### 2.12 *In situ* hybridization

For the expression analysis of *Cyp4g1* mRNA in the fat body, digoxigenin (DIG)-labelled sense and anti-sense probes (1001 bp) were synthesized by in vitro transcription with a DIG RNA Labelling Mix (Roche, Basel, Switzerland). The procedures from the dissection to hybridization were performed as described by Koto et al., 2019. The fat body was dissected from newly mated *P. lamellidens* queens in ant saline and fixed in 4% formaldehyde in phosphate buffer for 20 min at room temperature. The tissues were rehydrated by a decreasing series of methanol (75%, 50%, and 25%) and PTw (0.1% Tween 20 in 0.1 M PBS) solutions for 10 min each after being maintained overnight in methanol at - 20°C. The rehydrated tissues were digested with 20 μg/ml proteinase K for 3 min in PTw, and then 2 mg/ml glycine in PTw was added twice for 15 min to stop the digestion process. The tissues were post-fixed in PTw with 4% PFA for 20 min after being rinsed with PTw and then transferred to 50% PTw in a hybridization buffer (50% formamide, 5XSSC, 0.1% Tween 20, 1X Denhardt’s solution, 1 mg/ml tRNA [Roche, Basel, Switzerland], 50 μg/ml heparin and 0.1 mg/ml herring sperm DNA [Wako Pure Chemical Industries, Tokyo, Japan]) for 10 min. The prehybridization and hybridization processes were carried out by using hybridization buffer and RNA probes. Tissues were hybridized overnight with 100 ng labelled anti-sense or sense RNA probes at 60°C after a prehybridization was performed with hybridization buffer for 2 h at 60°C. When the hybridization process was complete, we transferred tissues into decreasing concentrations of hybridization buffer (75%, 50%, 25%) in 2X SSC for 30 min each at 60°C to wash and incubated the tissues with 2x SSC and 0.2X SSC for 30 min each. After treatment with DIG buffer I (100 mM Tris-HCl pH 7.5, 150 mM NaCl) for 5 min, the tissues were treated with 1.5% blocking reagent (Roche) for 1 h to prevent non-specific adsorption of antibody. We performed antigen-antibody reactions using the anti-DIG antibody that was conjugated with alkaline phosphatase (1:1000; Roche) in DIG buffer I for 1 h. The tissues were washed with DIG buffer I twice for 15 min and then transferred into DIG buffer III (100 mM Tris-HCl pH 9.5, 100 mM NaCl, 50 mM MgCl2, and 0.005% Tween 20) for 3 min. The staining was carried out using NBT/BCIP stock solution (Roche) for 1 h, and then stop solution (TE pH 8.0) was applied for 3 min. Finally, the stained tissues were washed with DIG buffer III twice for 10 min and PTw for 5 min. The tissues were mounted with 70% glycerol. The images of mounted tissues were acquired by the VHX-5000 system (Keyence).

## 3 Results

### 3.1 Induction of rubbing behaviour under laboratory conditions

To perform quantitative assessments, first, the rubbing behaviour of a newly mated *P. lamellidens* queen (the new queen) was re-enacted in a laboratory environment (Figure 1, Supplementary Movie 1). The rubbing behaviour experiment was carried out in a plaster-lined plastic case (5.0 × 4.5 × 2.5 cm), and the new queen interacted with the host workers in the case. To reduce the risk of counterattack from the host workers, the host workers were anaesthetized before being placed with the new queen (the detailed procedures are described in the Methods section). The rubbing behaviour observed in the field is characterized by the queens straddling the host worker and rubbing it with her legs (Figure 1A; Kohriba, 1963 & 1966; Kubota, 1974; Sakai, 1990, 1996, & 2000; Yano, 1911), and we succeeded in observing the same behaviour in the laboratory environment. In addition, this rubbing behaviour of the new queen was similar for both known host workers, *C. japonicus* (Japanese Ant Database Group, 2003; Kohriba, 1963) and *C. obscuripes* (Iwai et al., 2021) (Figures 1BC, Supplementary Movie 1).

### 3.2 The effect of rubbing behaviours on host workers

A behavioural test was performed to examine how rubbing behaviours changed the nestmate discrimination imposed by host workers. The behavioural test measured the actions of host workers towards the new *P. lamellidens* queen after three days of rubbing behaviour was observed on the host worker. At this time, the rubbing behaviour was performed on the worker from the same colony as that of the workers used for the behavioural test. When the newly mated *P. lamellidens* did not perform rubbing behaviours, *C. japonicus* workers recognized *P. lamellidens* as a non-nestmate and a high frequency of aggressive behaviour was observed (Figure 2A). On the other hand, *C. japonicus* workers did not attack the new queen that had exhibited rubbing behaviour (Figure 2A). This finding suggests that the rubbing behaviour enabled the newly mated queen of *P. lamellidens* to deceive the system of host nestmate recognition.

**Figure 2.**
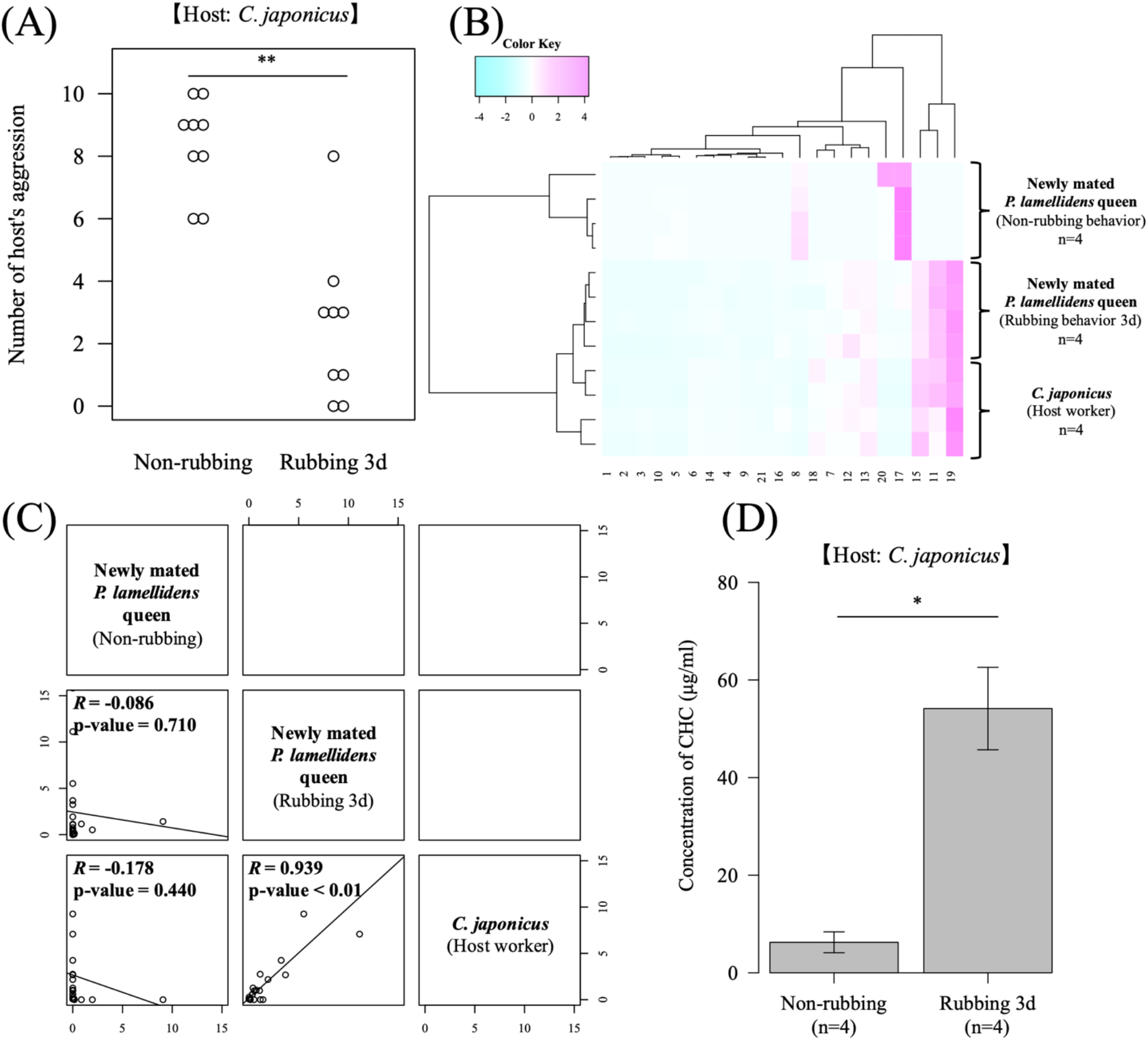
The CHC profile and effect of rubbing behaviour on the host worker reaction. We used *C. japonicus* as a host worker in these experiments. (A) The number of aggressive behaviours of host workers against newly mated *P. lamellidens* queens. These host workers were prepared with nine biological replicates, and they belonged to the same colony as that used in the rubbing behaviour experiments but with different individuals. Ten actions were observed against the newly mated *P. lamellidens* queen, and among these actions, “threaten”, “rush”, and “bite” were used to count the number of host aggressions. Wilcoxon signed rank test, **; significant difference: p value < 0.01. (B) Hierarchical clustering analysis of CHC profiles of newly mated *P. lamellidens* queens and host workers. This analysis was performed using the standardized value of the peak area that was converted to a Z score. The x-axis indicates the type of estimated hydrocarbons (see Supplementary Table 2). (C) Correlation analysis of CHC profiles of newly mated *P. lamellidens* queens and host workers. The correlation coefficient was calculated by the Pearson method. *R* indicates the correlation coefficient. Each plot shows the detected cuticular hydrocarbons. The x-axis and the y-axis show the area values of the standardized average peak area value. (D) The concentration of CHC in newly mated *P. lamellidens* queens. Mann–Whitney U test, *; significant difference: p value < 0.05, n=4.Error bar indicates standard error.

To determine how the rubbing behaviour altered the response of the host workers, a GC-MS analysis was conducted. The mass spectrometer measured the cuticular hydrocarbons (CHCs) on the body surface of new queens that had or had not exhibited the rubbing behaviour, as well as the host workers that were used as the rubbing target. As a result, the CHC profile of the newly mated *P. lamellidens* queens that had exhibited rubbing behaviour had clearly changed to one that closely resembling the host worker profile (Figures 2BC, Supplementary Figure 1, Supplementary Table 2, *R* > 0.7, p value < 0.01). Interestingly, the total amount of CHCs of the new queen increased due to the rubbing behaviour (Figure 2D). The profiles were also confirmed to change even when the target of rubbing behaviour was *C. obscuripes* (Supplementary Figure 2 and 3, Supplementary Table 3). These results suggest that the new queen can disguise her CHC profile and amount of CHSs by rubbing regardless of the host species to avoid host worker aggression.

### 3.3 Tracing assay with labelled CHCs

MS and behavioural tests demonstrated that the rubbing behaviour enables the host CHC profile to be disguised. However, whether this chemical disguise is implemented by chemical camouflage or chemical mimicry is unclear. Here, we focused on the result that the total amount of CHCs in the new queen was increased by the rubbing behaviour (Figure 2D). Therefore, whether this chemical disguise is implemented by direct deprivation from the body surface of the host worker (chemical camouflage) was verified by a tracing assay in which substances labelled with stable isotopes were applied. The tracking experiments were conducted using two types of hydrocarbons (n-triacontane-d62: C_30_D_62_, n-dotriacontane-d66: C_32_D_66_), in which the hydrogens of hydrocarbons with a chain length similar to those possessed by the host worker (*C. japonicus*; Supplementary Table 2) were replaced by stable isotopes. The efficiency of applying labelled substances to the ant body was confirmed with GC-MS measurements, and the labelled substances accounted for approximately 10-20% of the total CHC in the host worker (Supplementary Figure 4A). A similar ratio was also confirmed in the newly mated *P. lamellidens* queens, which exhibited rubbing behaviours (Supplementary Figure 4B). A new queen could perform rubbing behaviour on the labelled host worker for three days. By rubbing, the newly mated *P. lamellidens* queens acquired both the host CHC and the labelled substances on their cuticle from these host workers (Figures 3A–C, Supplementary Figure 4B and 5). In addition, since postpharyngeal glands are involved in the storage of hydrocarbons in ants (Soroker et al., 1994 & 1995), we analysed the hydrocarbon content in the postpharyngeal glands of the newly mated *P. lamellidens* queens. Host CHC and labelled substances were also detected in the postpharyngeal glands of newly mated *P. lamellidens* queens (Figure 3D, Supplementary Figure 6). These results suggest that the new queen directly obtains the host CHC through rubbing behaviour.

**Figure 3.**
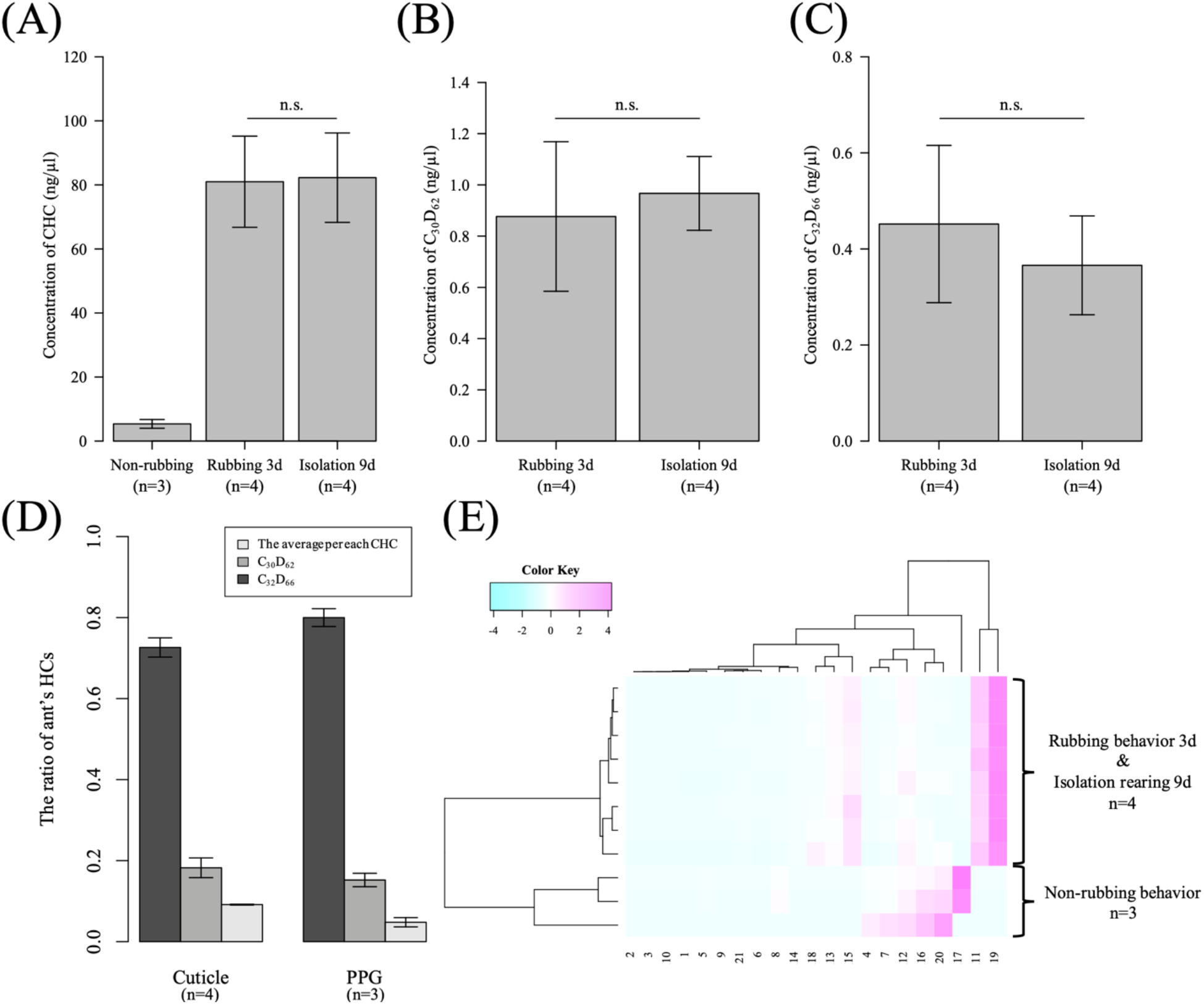
Comparison of CHC and labelled substances in the newly mated *P. lamellidens* queens. These experiments used *C. japonicus* as a host worker (A-E). (A) The concentration of CHC in the newly mated *P. lamellidens* queens (Mann–Whitney U test, n.s.; nonsignificant difference, n=4). (B&C) The concentration of labelled substances (n-triacontane-d62: C_30_D_62_, n-dotriacontane-d66: C_32_D_66_) in newly mated *P. lamellidens* queens (Mann–Whitney U test, n.s.; non-significant difference, n=4). (D) The ratio of hydrocarbons (the average per CHC, C_30_D_62_, C_32_D_66_) on the newly mated *P. lamellidens* queen cuticle and postpharyngeal glands (PPG) after rubbing behaviour. The average ratios of hydrocarbons identified in the cuticle and postpharyngeal glands were correlated (Pearson method, *R*=1, p value<0.01). (E) Hierarchical clustering analysis of the CHCs of newly mated *P. lamellidens* queens. This analysis was performed using the peak area value that was converted to a Z score. The number on the X-axis indicates the type of estimated hydrocarbon (see Supplementary Table 2). All error bars indicate the standard error.

**Figure 4.**
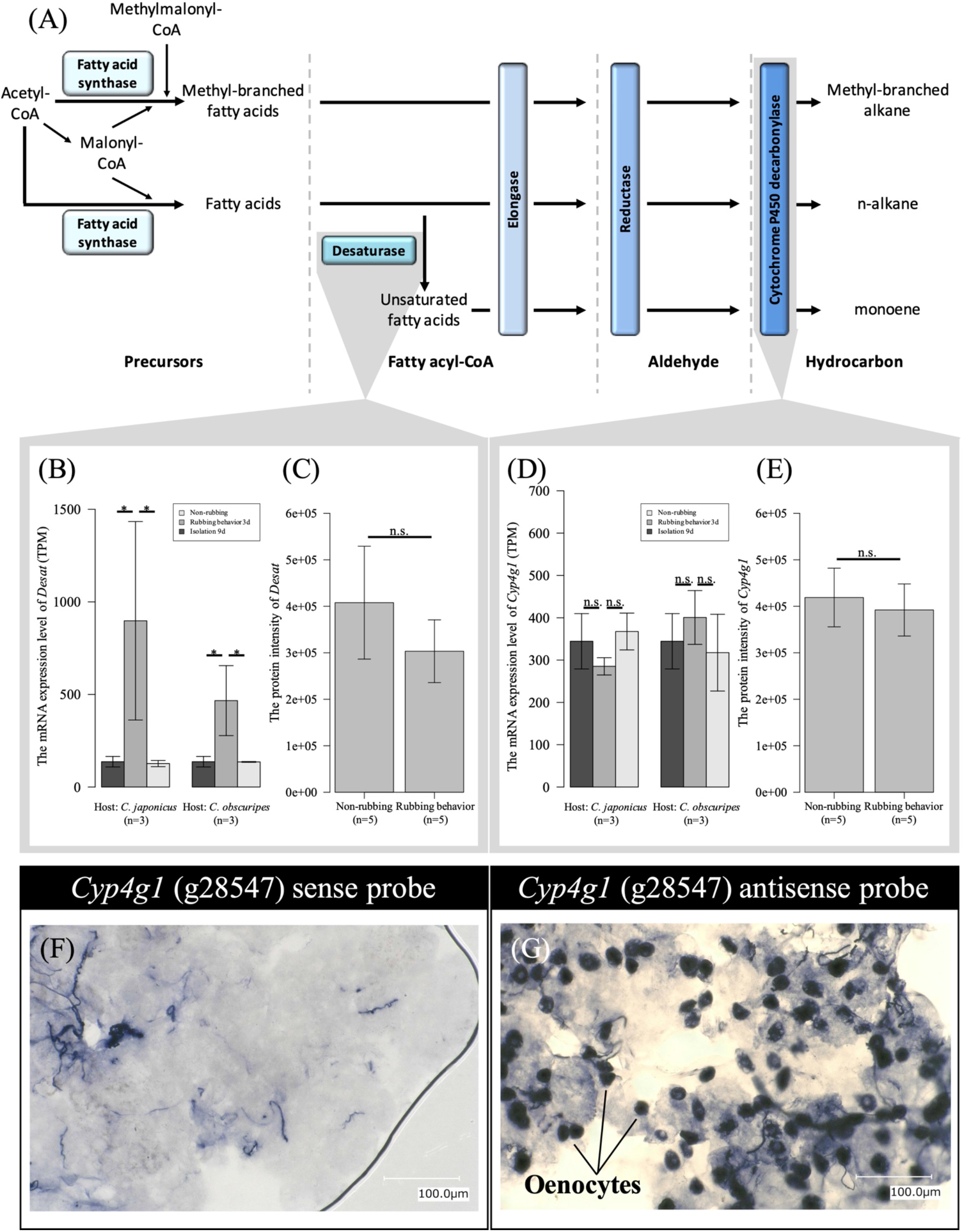
The expression levels of CHC synthesis-related genes in the whole abdomen. (A) The synthesis pathway of CHC (modified from Chung & Carroll, 2015). (B&D) The expression levels of desaturase (*Desat*): g34819.t1 in mRNA from whole abdomen by transcriptome (B) and protein from fat body by proteome (C). The expression levels of cytochrome P450 decarbonylase (*Cyp4g1*): g28547.t1 in mRNA from whole abdomen by transcriptome (D) and protein from fat body by proteome (E). In mRNA, *; Significant difference: FDR < 0.05, n.s.; non-significant difference, n=3. In protein, one-way ANOVA, n.s.; non-significant difference, n=5). (F&G) *In situ* hybridization using a *Cyp4g1* sense probe (F) and antisense probe (G) in the fat body. All error bars indicate the standard error.

To observe how long the new queen can retain the acquired CHC, including the labelled substances, the new queens were isolated for up to nine days after rubbing behaviour was observed against the labelled host worker, and the amount of CHC and CHC profile were measured. The newly mated *P. lamellidens* queen maintained the disguised CHC profile for at least nine days of isolation (Figure 3E, Supplementary Figures 7AB). Furthermore, the quantity of disguised CHC and labelled substances did not significantly increase or decrease during the nine days (Figures 3A–C, Supplementary Figure 7C).

### 3.4 Gene and protein expression profiling

The tracing assay with labelled CHC suggested that the chemical disguise of *P. lamellidens* may be based on a “chemical camouflage” method. We next examined another disguising strategy, “chemical mimicry”. Since chemical mimicry is a biosynthesis-mediated strategy, expression profiling was performed. A reference transcriptome of *P. lamellidens* was established by extracting the total RNA from the whole body of the larvae, worker, queen, abdomen of the newly mated queen, and fat body of the newly mated queen. cDNA was synthesized from the obtained total RNA and sequenced by an Illumina sequencer. Paired-end and single-end sequencing of 75–150 bp produced a total of 210 M reads (Supplementary Table 4). We assembled these reads using Trinity v2.8.5 and constructed the reference assembly of 23,523 coding regions with a BUSCO completeness of 96.6% (Supplementary Table 5).

The expression levels of the newly mated queen were analysed by RNA-seq with the *P. lamellidens* reference assembly, and these analyses were performed for queens without rubbing behaviour (non-rubbing behaviour) and queens with rubbing behaviour to the host workers for three days (rubbing 3d), or queens that were isolated from the hosts for nine days after rubbing behaviour was observed (isolation 9d). There are five CHC synthesis-related genes (Figure 4A), and these were expressed on the oenocytes that are contained in the abdomen fat body (Koto et al., 2019; Martins & Ramalho-Ortigao, 2012; Roma et al., 2006 & 2008; Thiele & Camargo-Mathias, 2003). Fatty acid synthase is involved in the biosynthesis of CHC precursors, desaturase and elongase in the construction of CHC varieties (it is involved in extending the chain length and inserting double bonds), and fatty acyl-CoA reductase and cytochrome P450 decarbonylase are involved in the final modification of CHC. The expression level of most of the genes that are related to CHC synthesis did not change among the non-rubbing, rubbing 3d, and isolation 9d groups, except for desaturase (*Desat*) (Figure 4, Supplementary Table 6-8). Desaturase (*Desat*) (g34819.t1), a CHC synthesis-related gene, was changed at the mRNA level but not at the protein level (Figures 4BC, Supplementary Figures 8AB, Supplementary Table 7 and 8). The expression level of cytochrome P450 decarbonylase (*Cyp4g1*), which is specifically expressed in the oenocyte (Koto et al., 2019), is particularly known to directly correlate with the amount of synthesized CHC. In this study, cytochrome P450 decarbonylase (*Cyp4g1*) (g28547.t1) was not detected as a differentially expressed gene (DEG) by transcriptome analysis (Figures 4D-G, Supplementary Figure 8CD, Supplementary Table 7 and 8). In addition, the expression pattern of cytochrome P450 decarbonylase (*Cyp4g1*) was examined by RT–qPCR analysis (Supplementary Figure 8D) and proteomics (Figure 4E).

## 4 Discussion

Rubbing behaviour was shown to disguise the host-like CHC profile and prevent the aggression of hosts, and this was demonstrated through replicating rubbing behaviour in the laboratory, testing the nestmate-recognition behaviour by hosts, and performing GC-MS analysis with the newly mated *P. lamellidens* queen against its host worker. In addition, the newly mated *P. lamellidens* queens acquired and maintained artificially-applied labelling substances from the host workers by performing rubbing behaviour, while gene expression profiling showed no change in the expression of cytochrome P450 decarbonylase (*Cyp4g1*), which is a CHC synthesis-related factor, during rubbing behaviour. These results suggest that the newly mated *P. lamellidens* queens directly acquire the hosts’ CHC through rubbing behaviour and deceive the technique of nestmate recognition by hosts through employing chemical disguises.

In a previous rearing study, a reduction in aggressive behaviour was observed in host workers towards the newly mated *P. lamellidens* queen after rubbing behaviour was performed (Kohriba, 1963). Our results demonstrate that the effect is due to chemical disguise (Figure 2) and quantitatively indicate that versatility promotes this disguise in both host species (*C. japonicus* and *C. obscuripes*), which exhibit different CHC profiles (Figures 2, Supplementary Figure 1-3, Supplementary Table 2 and 3). In addition, we revealed that the newly mated *P. lamellidens* queens possessed only a minute amount of CHCs before the rubbing behaviour (Figure 2D, Supplementary Table 2 and 3), and this quantity was significantly increased after the rubbing behaviour for *C. japonicus* (Figure 2D). Obtaining a new CHC is natural and simple when the baseline is limited, and this observation also supports the idea of chemical camouflage (acquirement of CHC). Since a low base quantity of CHC has been observed in other social parasitic ants, we expect it to be one of their general strategies (d’Ettorre & Errard, 1998; Johnson et al., 2001; Martin et al., 2007; Tsuneoka & Akino, 2012). The tracing assay with the labelled substances revealed that the newly mated *P. lamellidens* queen acquires labelled substances from its host workers by performing rubbing behaviours (Figures 3B–D, Supplementary Figure 4B, 5 and 6). This result also confirmed that the newly mated *P. lamellidens* queens directly obtained host CHCs through rubbing behaviours. Previous studies reported that ants exchange CHC through performing trophallaxis and grooming with nestmates, and through these behaviours, the CHC profile becomes standardized within the colony (Dahbi et al., 1999; Soroker et al., 1994 & 1995). The newly mated *P. lamellidens* queen also grooms the host while performing rubbing behaviours, and the queen probably acquires the CHC of hosts during this process (Kubota, 1974).

Isolation rearing revealed that the newly mated queen, which was isolated from its hosts for up to nine days, could maintain the disguised CHC profile (Figures 3A–C, Figure 3E, Supplementary Figure 7). This result suggests that the newly mated queen could maintain the amount and profile of the CHC that was acquired from the host. It is known that ants store CHC in the postpharyngeal glands in their heads (Soroker et al., 1994 & 1995). We also confirmed that the postpharyngeal glands of newly mated *P. lamellidens* queens contain host CHC and labelled substances from its labelled host workers after performing rubbing behaviours (Figure 3D, Supplementary Figure 6). The hydrocarbon profile found in the postpharyngeal glands was similar to that of the host CHC, and the labelled substances applied to the host were also detected in the postpharyngeal glands, suggesting that the newly mated *P. lamellidens* queen also supplies the acquired host CHC to its postpharyngeal glands. Most likely, the newly mated queen enables the disguised CHC profile to be maintained by storing the acquired CHC in the postpharyngeal glands.

Expression analysis was performed to verify the chemical mimicry, and except for the mRNA of desaturase (*Desat*) (but not the protein level), no changes were observed in the expression of genes in the pathways related to CHC synthesis before and after rubbing behaviour (Figure 4, Supplementary Figure 8, Supplementary Table 6-8). This result suggests that the biosynthesis of CHC, i.e., chemical mimicry, does not occur in the newly mated *P. lamellidens* queens. Among the CHC synthesis-related genes, the expression of cytochrome P450 decarbonylase (*Cyp4g1*) (Koto et al., 2019), which is responsible for the final modification of CHC precursors and has an expressional level that correlates with the amount of CHC synthesized, was not detected at the mRNA or protein level (Figures 4D–G, Supplementary Figures 8CD, Supplementary Table 6-8). Since the expression of cytochrome P450 decarbonylase (*Cyp4g1*) did not change despite the apparent increase in the amount of CHC after rubbing behaviour, we expected that no new biosynthesis of CHC would occur, at least during the rubbing behaviour. The expression level of Desaturase (*Desat*), which is involved in the construction of CHC variants (Chung & Carroll, 2015), changed at the mRNA level but not at the protein level (Figures 4BC, Figures 8AB, Supplementary Table 6-8). Based on the above observations, we expected that the newly mated *P. lamellidens* queen would achieve chemical disguise by performing chemical camouflage (acquiring CHC) rather than chemical mimicry (CHC biosynthesis) through rubbing behaviour.

Most previous studies that were aimed at elucidating the mechanisms of chemical disguise verified the persistence of disguised CHC by isolating myrmecophile from its host (Akino et al., 1996; Scarparo et al., 2019; Vander Meer & Wojcik, 1982; von Beeren et al., 2011 & 2012). These studies revealed whether the CHC profile returns to its pre-disguise state when myrmecophiles are isolated from their hosts. The researchers believed that chemical disguise is carried out by chemical camouflage in the former case and chemical mimicry in the latter case. However, the results of isolation rearing and the tracing assay of labelled substances in this study showed that the newly mated *P. lamellidens* queen maintained CHC disguised by chemical camouflage for a certain period even in the absence of the host. These results suggest that even if the parasite can maintain a disguised CHC for a long time by isolation rearing, it may not be due to the new biosynthesis of CHC (chemical mimicry). To elucidate the mechanisms of chemical disguise, in addition to performing isolation rearing, it is necessary to construct an appropriate experimental system that corresponds to the parasite.

We used a combination of experimental techniques to reveal the chemical disguise strategy of the newly mated *P. lamellidens* queen in the early stages of social parasitism. In addition, we believe that this study succeeded in elucidating some of the previously obscure mechanisms that are involved in the chemical disguise of socially parasitic ants.

## 6 Author Contributions

HI and NK designed the project. HI collected ant specimens and performed transcriptome analysis. HI and MM conducted proteome analysis. The first draft of the manuscript was written by HI, and all authors commented on previous versions of the manuscript. NK managed the experimental environment. MT and KA managed the resources.

## 8 Acknowledgments

The authors thank Akiko Koto for supporting the *in situ* hybridization; Toshiharu Akino for providing extraction method of CHC; Masataka Wakayama, Noriko Fukuda, and Noriko Kagata for the supporting GC-MS measurement; and Yuki Takai for technical support in RNA sequencing. The authors also thank Daiki D Horikawa, Keizo Takasuka, Takahiro Masuda, Konosuke Ii, Yu Kurihara, Sora Ishikawa, Phillip Yamamoto, and Tomoki Takeda for their critical suggestions.

## 9 Data Availability Statement

The datasets of transcriptome and proteome for this study can be found in SRA [accession number: SUB11231698] and JPOST [accession number: JPST001552.2]. The transcriptome assembly can also be found in TSA [accession number: GJVV01000000 (the sequences less than 200 bp in registered assembly were removed)].

