## Supplementary_Material for "Molecular evidence of chemical disguise by the socially parasitic spiny ant *Polyrhachis lamellidens* (Hymenoptera: Formicidae) when invading a host colony"

### 1. Supplementary Tables

#### Supplementary Table 1. The list of primers for RT–qPCR

| Gene name | Primer type | Sequence (5' to 3') |
|---|---|---|
| *Gapdh1* (g57081.t1) | Forward | CGTATTGGCCGTCTTGTACTG |
| | Reverse | CGGCTTTGACTTCTCCCTTG |
| *Actin5C* (g5939.t1) | Forward | TCGCGACATCAAGGAAAAGC |
| | Reverse | ACAGCGGAATCGTTCGTTAC |
| *Desat* (g34819.t1) | Forward | GGAAGTCAAGGAAAAGGGCAAG |
| | Reverse | TCCAACACACAACCGGAATG |
| *Cyp4g1* (g28547.t1) | Forward | TGCGCCTTAGGAAATCATCAG |
| | Reverse | ACGGTACTGGTGGGAAAAGTC |

**Supplementary Table 2. The list of estimated CHCs from newly mated *P. lamellidens* queens and *C. japonicus*.** +++/- indicates CHCs that were commonly detected in all ant samples or not detected at all. CHCs detected in more or less than half of ant samples are represented by ++ or +. ECL: equivalent chain length of n-alkane. The number of CHCs was counted from ant samples which *C. japonicus* was used in the experiment.

| Peak no. | ECL | CHC | *P. lamellidens* queen Non-rubbing behaviour | *P. lamellidens* queen (Isolated after rubbing behaviour) | *C. japonicus* worker |
|---|---|---|---|---|---|
| 1 | | $C_{23:1}$ | - | + | +++ |
| 2 | 23 | $nC_{23}$ | - | +++ | +++ |
| 3 | | $C_{25:1}$ | - | ++ | +++ |
| 4 | 25 | $nC_{25}$ | ++ | +++ | +++ |
| 5 | | 7,12-dime$C_{25}$ | ++ | ++ | - |
| 6 | | $C_{26:1}$ | - | +++ | +++ |
| 7 | 26 | $nC_{26}$ | ++ | +++ | +++ |
| 8 | | 5,10-dime$C_{26}$ | ++ | + | - |
| 9 | | 5,7,12- and 7,9,12-trime$C_{25}$ | - | +++ | +++ |
| 10 | | Unknown alkane | + | ++ | - |
| 11 | | $C_{27:1}$ | - | +++ | +++ |
| 12 | 27 | $nC_{27}$ | ++ | ++ | +++ |
| 13 | | 13-me$C_{27}$ | - | +++ | +++ |
| 14 | | 5-me$C_{27}$ | - | +++ | +++ |
| 15 | | 7,15-dime$C_{27}$ | - | +++ | +++ |
| 16 | 28 | $nC_{28}$ | ++ | ++ | +++ |
| 17 | | 5,10,15- trime$C_{27}$ | ++ | ++ | - |
| 18 | | 5,7,12-trime$C_{27}$ | - | ++ | +++ |
| 19 | | $C_{29:1}$ | - | +++ | +++ |
| 20 | 29 | $nC_{29}$ | ++ | ++ | - |
| 21 | | 13-me$C_{29}$ | - | +++ | +++ |

**Supplementary Table 3. The list of estimated CHCs from newly mated *P. lamellidens* queens and *C. obscuripes*.** +++/- indicates CHCs that were commonly detected in all ant samples or not detected at all. CHCs detected in more or less than half of ant samples are represented by ++ or +. ECL: equivalent chain length of n-alkane. The number of CHCs was counted from ant samples which *C. obscuripes* was used in the experiment.

| Peak no. | ECL | CHC | *P. lamellidens* queen Non-rubbing behaviour | *P. lamellidens* queen (Isolated after rubbing behaviour | *C. obscuripes* worker |
|---|---|---|---|---|---|
| 2 | 23 | $nC_{23}$ | - | ++ | +++ |
| 22 | 24 | $nC_{24}$ | - | ++ | +++ |
| 4 | 25 | $nC_{25}$ | - | +++ | +++ |
| 7 | 26 | $nC_{26}$ | - | +++ | +++ |
| 8 | | 5,10-dime$C_{26}$ | + | +++ | - |
| 23 | | 2-me$C_{26}$ | - | +++ | +++ |
| 11 | | $C_{27:1}$ | - | +++ | +++ |
| 12 | 27 | $nC_{27}$ | +++ | +++ | +++ |
| 24 | | 11-me$C_{27}$ | - | +++ | +++ |
| 25 | | $C_{28:1}$ | - | +++ | +++ |
| 16 | 28 | $nC_{28}$ | - | +++ | +++ |
| 17 | | 5,10,15- trime$C_{27}$ | +++ | +++ | - |
| 26 | | 3-me$C_{28}$ | - | +++ | +++ |
| 19 | | $C_{29:1}$ | - | +++ | +++ |
| 20 | 29 | $nC_{29}$ | +++ | +++ | +++ |
| 27 | | Unknown alkane | - | +++ | +++ |
| 28 | | 9,13-dime$C_{29}$ | - | +++ | +++ |
| 29 | | $C_{30:1}$ | - | +++ | +++ |
| 30 | 30 | $nC_{30}$ | - | +++ | +++ |
| 31 | | 12-me$C_{30}$ | - | +++ | +++ |
| 32 | | Unknown alkane | - | +++ | +++ |
| 33 | | $C_{31:1}$ | - | +++ | +++ |
| 34 | 31 | $nC_{31}$ | - | +++ | +++ |
| 35 | | Unknown alkane | - | +++ | +++ |
| 36 | | 11,16-dime$C_{31}$ | - | +++ | +++ |
| 37 | | Unknown alkane | - | +++ | +++ |
| 38 | | 10,16-dime$C_{32}$ | - | +++ | +++ |
| 39 | | 11,17-dime$C_{33}$ | - | +++ | +++ |
| 40 | | Unknown alkane | - | +++ | +++ |
| 41 | | Unknown alkane | - | +++ | +++ |

**Supplementary Table 4. The list of RNA-Seq data from *P. lamellidens* obtained in this work.** The BioProject ID of these data is SUB11233824. *Each data was used to transcriptome assembly.

| Sample type | Body part | Description | Sequence type | Accession No. | |
|---|---|---|---|---|---|
| Larva | Whole body | The pool of five larvae | Paired-end | SRR18532324 | * |
| Worker | Whole body | | Paired-end | SRR18532323 | * |
| Queen | Whole body | | Paired-end | SRR18532312 | * |
| Newly mated queen | Abdomen | No rubbing behaviour | Single-end | SRR18532303 | * |
| Newly mated queen | Abdomen | No rubbing behaviour | Single-end | SRR18532302 | * |
| Newly mated queen | Abdomen | No rubbing behaviour | Single-end | SRR18532301 | * |
| Newly mated queen | Abdomen | Rubbing to the *C. japonicus* workers for 3 days | Single-end | SRR18532300 | * |
| Newly mated queen | Abdomen | Rubbing to the *C. japonicus* workers for 3 days | Single-end | SRR18532299 | * |
| Newly mated queen | Abdomen | Rubbing to the *C. japonicus* workers for 3 days | Single-end | SRR18532298 | * |
| Newly mated queen | Abdomen | Rubbing to the *C. obscuripes* workers for 3 days | Single-end | SRR18532297 | * |
| Newly mated queen | Abdomen | Rubbing to the *C. obscuripes* workers for 3 days | Single-end | SRR18532322 | * |
| Newly mated queen | Abdomen | Rubbing to the *C. obscuripes* workers for 3 days | Single-end | SRR18532321 | * |
| Newly mated queen | Abdomen | Isolated for 9 days after the rubbing behaviour to the *C. japonicus* workers | Single-end | SRR18532320 | * |
| Newly mated queen | Abdomen | Isolated for 9 days after the rubbing behaviour to the *C. japonicus* workers | Single-end | SRR18532319 | * |
| Newly mated queen | Abdomen | Isolated for 9 days after the rubbing behaviour to the *C. japonicus* workers | Single-end | SRR18532318 | * |
| Newly mated queen | Abdomen | Isolated for 9 days after the rubbing behaviour to the *C. obscuripes* workers | Single-end | SRR18532317 | * |
| Newly mated queen | Abdomen | Isolated for 9 days after the rubbing behaviour to the *C. obscuripes* workers | Single-end | SRR18532316 | * |
| Newly mated queen | Abdomen | Isolated for 9 days after the rubbing behaviour to the *C. obscuripes* workers | Single-end | SRR18532315 | * |
| Newly mated queen | Fat body | No rubbing behaviour | Single-end | SRR18532314 | |
| Newly mated queen | Fat body | No rubbing behaviour | Single-end | SRR18532313 | |
| Newly mated queen | Fat body | No rubbing behaviour | Single-end | SRR18532311 | |
| Newly mated queen | Fat body | No rubbing behaviour | Single-end | SRR18532310 | |
| Newly mated queen | Fat body | Rubbing to the *C. japonicus* workers for 3 days | Single-end | SRR18532309 | |
| Newly mated queen | Fat body | Rubbing to the *C. japonicus* workers for 3 days | Single-end | SRR18532308 | |
| Newly mated queen | Fat body | Rubbing to the *C. japonicus* workers for 3 days | Single-end | SRR18532307 | |
| Newly mated queen | Fat body | Rubbing to the *C. japonicus* workers for 3 days | Single-end | SRR18532306 | |
| Newly mated queen | Fat body | Rubbing to the *C. japonicus* workers for 3 days | Single-end | SRR18532305 | |
| Newly mated queen | Fat body | Rubbing to the *C. japonicus* workers for 3 days | Single-end | SRR18532304 | |

**Supplementary Table 5. Assembly statistics of the *P. lamellidens* transcriptome.**

| | ***P. lamellidens*** |
|---|---|
| Number of predicted genes | 23,523 |
| Average scaffold length | 1,187 bp |
| Total scaffold length | 27,914,451 bp |
| Longest scaffold | 40,884 bp |
| Scaffold N50 | 2,187 bp |
| BUSCO (Arthropoda) | 96.6 % |

**Supplementary Table 6. The number of differentially expressed genes (DEGs) in the abdomen (FDR < 5%).**

| | ***C. japonicus*** | ***C. obscuripes*** | **Common in both** |
|---|---|---|---|
| Non-rubbing vs. Rubbing 3d | 65 | 103 | 20 |
| Rubbing 3d vs. Isolation 9d | 807 | 247 | 74 |

**Supplementary Table 7. The candidate CHC synthesis-related genes in transcriptome assembly.**

| Gene name | UniProt ID | Gene ID | Identity of BLAST (%) |
|---|---|---|---|
| Fatty acid synthase (*FASN2*) | Q9VQL6_DROME, M9PB21_DROME | g30275.t1 | 49.461-49.482 |
| Elongase (*eloF*) | ELOF_DROME | g39123.t1, g56534.t1, g36733.t1, g21655.t1, g23416.t1, g23415.t1, g24854.t1 | 34.000-38.554 |
| Desaturase (*Desat1, Desat2, desatF*) | Q7K4Y0_DROME, Q9VG68_DROME, A7DZ97_DROME | g34819.t1, g11236.t1, g11238.t1, g53824.t1, g49648.t1, g53826.t1 | 37.829-69.006 |
| Fatty acyl-CoA reductase (*Sgp*, *anon-WO0140519.58*, *anon-WO0140519.58*, *Dmel*＼*CG4020*) | Q9VLJ7_DROME, Q9VG86_DROME, Q9VES6_DROME, Q9W459_DROME | g58736.t1, g43597.t1, g45448.t1, g45454.t1, g61926.t1, g24041.t1, g56854.t1, g56866.t1, g22537.t1, g24040.t1, g31109.t1, g24045.t1, g50718.t1, g50719.t1, g15289.t1, g18683.t1 | 27.105-41.628 |
| Cytochrome P450 decarbonylase (*Cyp4g1*) | CP4G1_DROME | g28547.t1, g28546.t1, g57012.t1, g20324.t1, g12257.t1, g37042.t1, g20315.t1, g40110.t1, g48658.t1, g48660.t1, g51446.t1, g64826.t1, g20314.t1, g598.t1, g42993.t1 | 24.416-53.802 |

**Supplementary Table 8. The annotated differentially expressed genes (DEGs) (FDR < 5%).** The predicted gene names of the DEGs that were commonly observed regardless of the host species. The logFC is written for the comparison group in which changes in expression were observed. *One candidate for cytochrome P450 decarbonylase (*Cyp4g1*), g57012.t1, was detected by RNA-Seq in the whole abdomen. However, the candidate was omitted from the discussion in the text because its pattern of expression was different in the fat body.

| **Gene ID** | **Gene name** | **Top hit accession ID** | **The logFC of DEGs** | |
|---|---|---|---|---|
| | | | **Non-rubbing vs. Rubbing 3d** | **Rubbing 3d vs. Isolation 9d** |
| g587.t1 | No hit | No hit | 1.38, 1.34 | - |
| g994.t1 | No hit | No hit | -5.88, -5.87 | - |
| g2502.t1 | Lysosomal alpha-mannosidase isoform X1 | XP_025265136.1 | - | 2.11, 1.43 |
| g3009.t1 | Fatty acid-binding protein homologue 7-like | XP_029664361.1 | - | -1.21, -1.88 |
| g3391.t1 | Uncharacterized protein | XP_029666367.1 | - | -1.68, -1.13 |
| g4461.t1 | Facilitated trehalose transporter Tret1-2 homologue | XP_011263555.1 | - | -1.92, -3.06 |
| g5567.t1 | Uncharacterized protein | XP_011262925.2 | - | -5.69, 6.34 |
| g6106.t1 | Ubiquitin-conjugating enzyme E2 G2 | XP_011636099.1 | 2.59, 2.22 | - |
| g7723.t1 | Ribose-phosphate pyrophosphokinase 1 isoform X1 | XP_029172996.1 | - | 2.37, 1.96 |
| g7979.t1 | Hypothetical protein | EFN70423.1 | - | -1.72, -1.21 |
| g8968.t1 | Sodium- and chloride-dependent betaine transporter | XP_011260796.1 | - | -1.59, -2.23 |
| g9111.t1 | Fatty acid synthase-like | XP_011264667.2 | 1.84, 1.63 | - |
| g9808.t1 | No hit | No hit | 3.83, -7.67 | - |
| g10769.t1 | No hit | No hit | - | 0.71, 0.94 |
| g11011.t1 | Uncharacterized protein | XP_029665965.1 | 1.52, 1.32 | - |
| g11948.t1 | Spidroin-2-like isoform X1 | XP_029661708.1 | - | -1.65, -1.95 |
| g13735.t1 | Haemocyte protein-glutamine gamma-glutamyltransferase | XP_011266432.1 | - | -0.87, -1.26 |
| g15692.t1 | Metabotropic glutamate receptor isoform X1 | XP_025266613.1 | - | -2.41, -2.82 |
| g16463.t1 | Leucine-rich repeat-containing protein 15 | XP_011253803.1 | - | -0.88, -1.40 |
| g16588.t1 | Fibrillin-1 | XP_025262768.1 | - | -1.18, -1.58 |
| g19442.t1 | Trifunctional purine biosynthetic protein adenosine-3 | XP_025268113.1 | - | 1.50, 1.91 |
| g20432.t1 | PREDICTED: uncharacterized protein | XP_012225306.1 | - | -1.46, -0.93 |

| | | | | |
|---|---|---|---|---|
| g22437.t1 | Arylsulfatase B isoform X1 | XP_025270578.1 | - | -0.92, -1.45 |
| g23508.t1 | Prostatic acid phosphatase isoform X2 | XP_011255200.1 | - | -1.33, -1.20 |
| g24953.t1 | No hit | No hit | - | -8.32, 8.31 |
| g25803.t1 | No hit | No hit | - | 2.67, 3.49 |
| g26264.t1 | Multidrug resistance protein homologue 49 isoform X1 | XP_011265189.1 | - | -1.07, -1.09 |
| g26732.t1 | Solute carrier family 2, facilitated glucose transporter member 1 isoform X2 | XP_011261955.2 | - | 1.16, 1.18 |
| g30275.t1 | Fatty acid synthase-like | XP_025270749.1 | - | -2.07, -1.23 |
| g31112.t1 | Scavenger receptor class B member 1 | XP_011253015.1 | - | -1.67, -2.06 |
| g33203.t1 | Uncharacterized protein | XP_011254770.1 | - | -1.72, -1.47 |
| g34129.t1 | No hit | No hit | -2.56, -7.27 | - |
| g34819.t1 | Acyl-CoA Delta (11) desaturase | XP_025264947.1 | 2.63, 1.73 | -2.85, -1.93 |
| g35207.t1 | WEB family protein At4g27595, chloroplastic | XP_025268350.1 | 2.80, 1.62 | - |
| g35887.t1 | Solute carrier organic anion transporter family member 2A1 | XP_011255071.1 | - | -1.38, -1.21 |
| g35963.t1 | Hypothetical protein | EFN74322.1 | - | 2.66, 1.58 |
| g35964.t1 | Hypothetical protein | EFN74322.1 | - | 2.30, 1.78 |
| g38487.t1 | Intraflagellar transport protein 43-like protein | KMQ94321.1 | - | 1.14, 1.41 |
| g38783.t1 | MFS-type transporter SLC18B1 | XP_011265521.1 | 0.53, 0.61 | -0.87, -1.51 |
| g39515.t1 | Calexcitin-2 | XP_011256346.1 | - | -1.44, -2.65 |
| g39681.t1 | Sodium-coupled monocarboxylate transporter 1-like | XP_025261690.1 | - | -1.12, -1.68 |
| g40149.t1 | NPC intracellular cholesterol transporter 2-like | XP_011258918.1 | - | -3.96, -1.94 |
| g40405.t1 | Uncharacterized protein | XP_029668922.1 | - | -1.09, -1.38 |
| g46969.t1 | Uncharacterized protein | XP_011264753.1 | - | -0.83, -1.19 |
| g47732.t1 | Lipid storage droplets surface-binding protein 1 | EFN73399.1 | - | -1.00, -1.05 |
| g49055.t1 | Glutamic acid-rich protein-like | XP_011259581.1 | - | 1.20, 0.96 |
| g49288.t1 | Pheromone-binding protein Gp-9-like isoform X2 | XP_029660118.1 | - | -3.11, -3.34 |
| g50866.t1 | Histone H2B-like | XP_011632649.1 | - | -1.71, -1.74 |
| g50909.t1 | Arylphorin subunit alpha-like | XP_029170871.1 | - | 3.02, -3.88 |
| g51070.t1 | No hit | No hit | - | -2.40, -8.86 |

| | | | | |
|---|---|---|---|---|
| g51590.t1 | No hit | No hit | - | 2.18, 4.47 |
| g51996.t1 | Venom acid phosphatase Acph-1-like | XP_025271241.1 | - | -1.27, -1.59 |
| g52645.t1 | Probable ascorbate-specific transmembrane electron transporter 1 | XP_025266889.1 | - | -1.81, -3.55 |
| g52952.t1 | Mitochondrial pyruvate carrier 1-like | XP_029665361.1 | - | -1.37, -1.17 |
| g53082.t1 | Sodium-coupled monocarboxylate transporter 1 isoform X1 | XP_025271097.1 | - | -1.29, -1.13 |
| g54876.t1 | L-lactate dehydrogenase | XP_011265340.1 | - | -1.10, -1.29 |
| g56481.t1 | Haemolymph lipopolysaccharide-binding protein-like | XP_011253802.2 | 2.16, 2.23 | -1.66, -2.43 |
| *g57012.t1 | Cytochrome P450 4g15 | XP_011267879.1 | 1.48, 1.54 | -1.28, -1.85 |
| g57346.t1 | Translation initiation factor if-2 | KMQ96767.1 | - | -1.42, -1.66 |
| g57976.t1 | No hit | No hit | - | -5.96, -6.55 |
| g58186.t1 | Feline leukaemia virus subgroup C receptor-related protein 2-like | XP_025261827.1 | - | -1.02, -1.05 |
| g58387.t1 | Inositol monophosphatase 1 | XP_011257136.1 | - | -1.54, -1.36 |
| g58775.t1 | Maltase 1-like | XP_025263040.1 | - | -2.39, -4.58 |
| g59001.t1 | Cytosolic 10-formyltetrahydrofolate dehydrogenase | XP_011251043.2 | - | -2.11, -2.49 |
| g59329.t1 | Uncharacterized protein | XP_025266319.1 | - | 1.29, -3.44 |
| g60423.t1 | No hit | No hit | - | -6.32, -4.87 |
| g60655.t1 | No hit | No hit | - | -2.50, -3.94 |
| g61485.t1 | Synaptic vesicle glycoprotein 2B | XP_011254928.1 | - | -0.98, -1.21 |
| g61864.t1 | Phospholipase B1, membrane-associated | XP_011257663.3 | 2.51, 2.02 | - |
| g62475.t1 | No hit | No hit | - | 10.03, 6.56 |
| g62621.t1 | Defensin | XP_025262447.1 | 7.13, 7.38 | -6.52, -8.17 |
| g63022.t1 | No hit | No hit | - | 1.31, 1.05 |
| g63075.t1 | No hit | No hit | - | 1.03, 1.03 |
| g63707.t1 | Uncharacterized protein | XP_025268004.1 | - | -6.57, -8.46 |
| g63709.t1 | Hymenoptaecin | KMQ95866.1 | 6.69, 7.80 | -7.72, -8.92 |
| g63711.t1 | Uncharacterized protein | XP_025268004.1 | 6.57, 7.74 | -7.44, -8.97 |
| g63714.t1 | Uncharacterized protein | XP_025268004.1 | 6.32, 7.69 | -7.34, -8.98 |
| g63715.t1 | Uncharacterized protein | XP_025268004.1 | 6.52, 7.96 | -7.61, -9.11 |

| | | | | |
|---|---|---|---|---|
| g64630.t1 | Cuticle protein 2-like | XP_025269546.1 | - | -1.60, -1.86 |
| g64805.t1 | Abnormal long morphology protein 1-like | XP_029162544.1 | - | -2.57, 3.39 |
| g65054.t1 | No hit | No hit | -6.06, -6.04 | - |
| g65360.t1 | No hit | No hit | - | -1.81, -2.89 |
| g67717.t1 | Facilitated trehalose transporter Tret1 | XP_025268384.1 | - | -0.98, -1.28 |
| g67726.t1 | Mediator of RNA polymerase II transcription subunit 12 | XP_025264718.1 | -1.80, 4.11 | - |
| g69521.t1 | No hit | No hit | 4.63, 3.50 | - |
| g69762.t1 | Larval cuticle protein A2B-like | XP_011260338.1 | - | -1.39, -1.74 |

### 2. Supplementary Figures

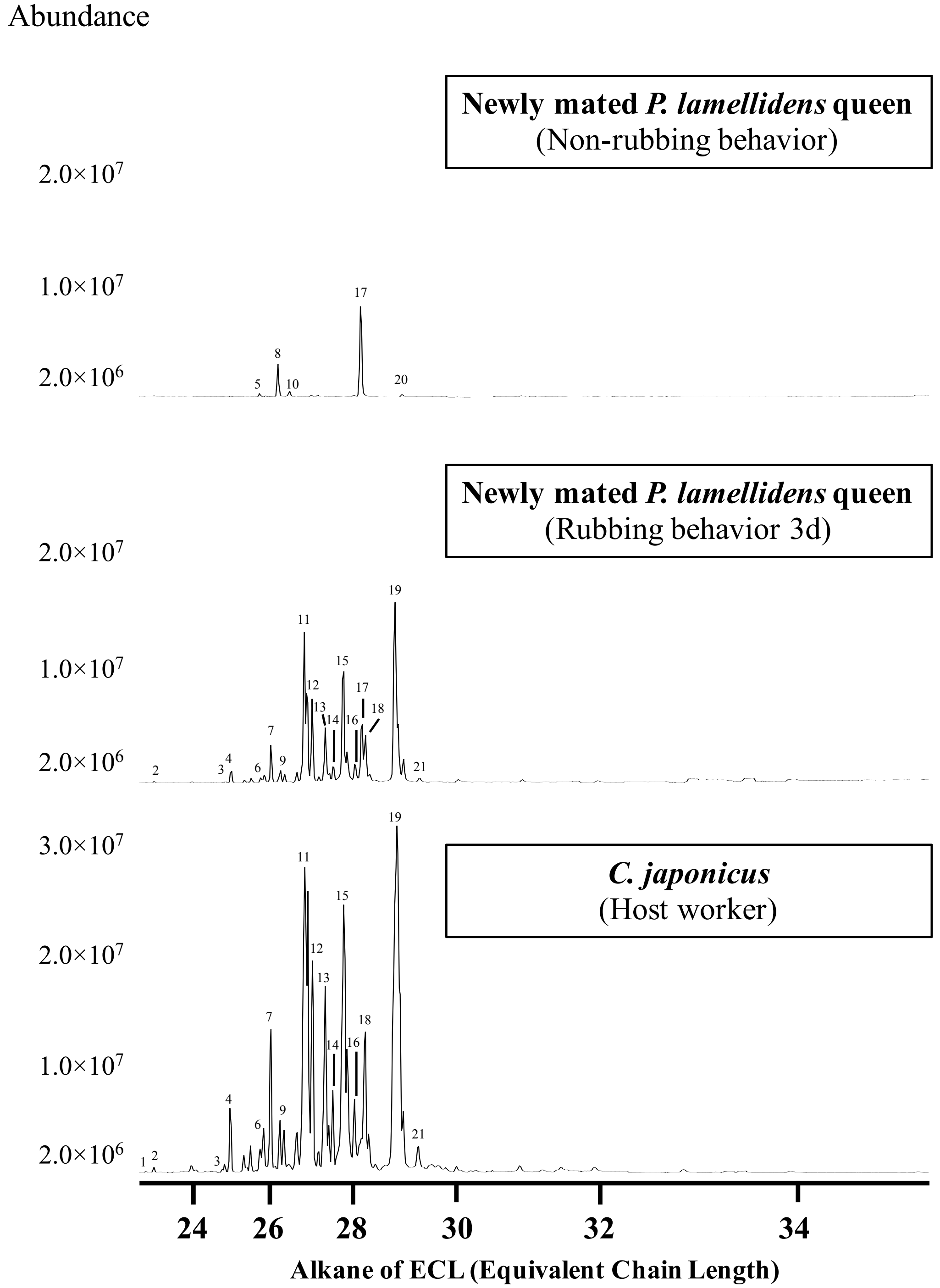


**Supplementary Figure 1. Examples of CHC chromatograms of newly mated *P. lamellidens* queens and their host (*C. japonicus*).** The number assigned to each peak indicates the type of estimated hydrocarbon (see Supplementary Table 2).

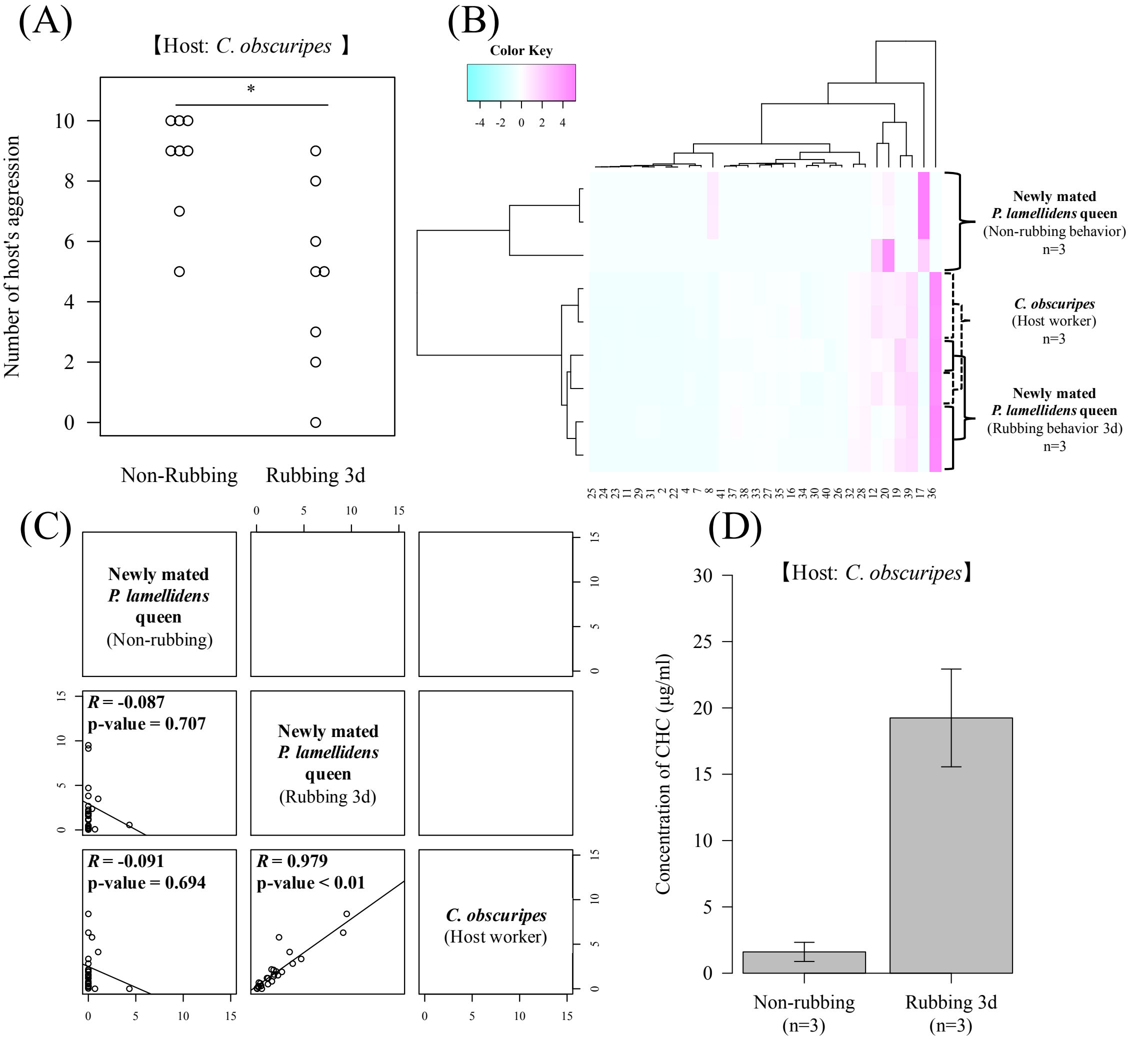


**Supplementary Figure 2. The CHC profile for the effect of rubbing behaviour on the host worker reaction.** We used *C. obscuripes* as a host worker in these experiments. (A) The number of incidences of aggressive behaviour by host workers towards newly mated *P. lamellidens* queens. These host workers were prepared with eight biological replicates, and they belonged to the same colony that was used in the rubbing behaviour experiments but the individual host workers were different. Ten actions by the host workers to the newly mated *P. lamellidens* queen were observed, and among these actions, “threaten”, “rush”, and “bite” were used to count the incidences of host aggression towards the queens. Wilcoxon signed rank test, *; significant difference: p value < 0.05. (B) Hierarchical clustering analysis of the CHC profiles of newly mated *P. lamellidens* queens and host workers. This analysis was performed using the standardized value of peak area, which was converted to the Z score. The x-axis indicates the type of estimated hydrocarbons (see Supplementary Table 3). (C) Correlation analysis of the CHC profiles of newly mated *P. lamellidens* queens and host workers. The correlation coefficient was calculated by the Pearson method. *R* indicates the correlation coefficient. Each plot shows the cuticular hydrocarbons that were detected. The x-axis and the y-axis show the area values of the standardized average peak area value. (D) The concentration of CHC in newly mated *P. lamellidens* queens. n=3. Error bar indicates standard error.

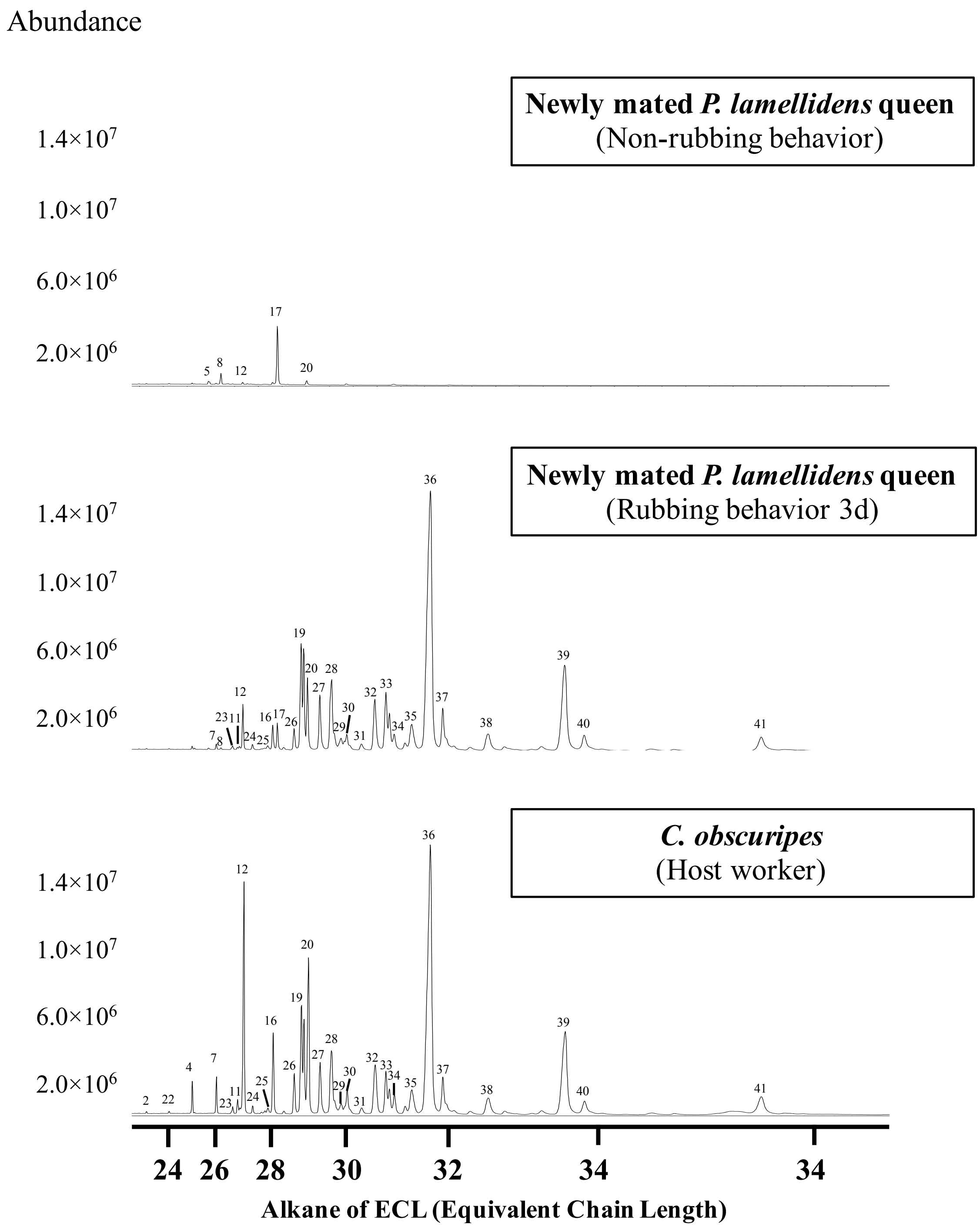


**Supplementary Figure 3. Examples of the CHC chromatograms of newly mated *P. lamellidens* queens and their hosts (*C. obscuripes*).** The number assigned to each peak indicates the type of estimated hydrocarbon (see Supplementary Table 3).

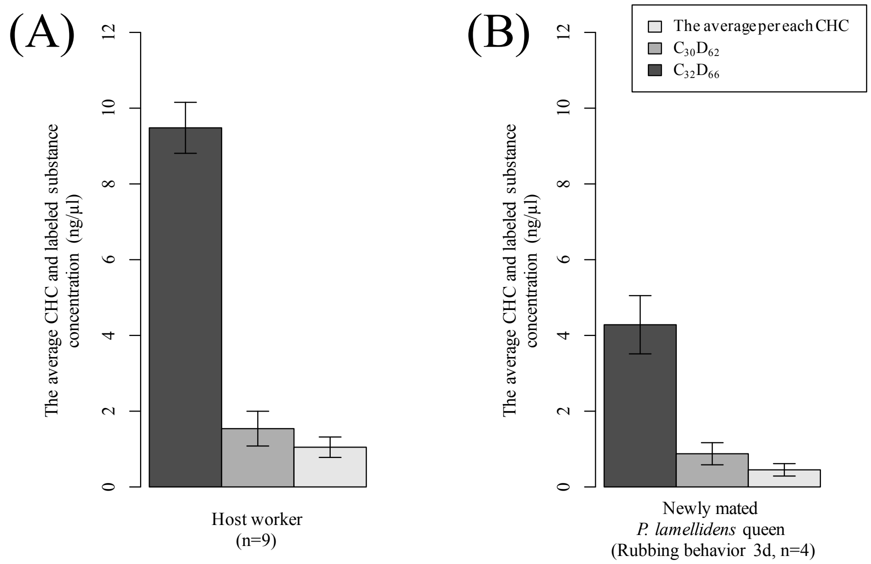


**Supplementary Figure 4. The concentration of CHC and labelled substances in newly mated *P. lamellidens* queens and host workers.** The concentration of labelled substances (n-triacontane-d62: $C_{30}D_{62}$, n-dotriacontane-d66: $C_{32}D_{66}$) and the average concentration per CHC of the host workers (*C. japonicus*) (A) and newly mated *P. lamellidens* queens (B). The host workers remained separated from the newly mated *P. lamellidens* queens for 1-3 days after the labelled substances were applied. For three days, the newly mated *P. lamellidens* queens exhibited rubbing behaviours towards a labelled host worker. All error bars indicate the standard error.

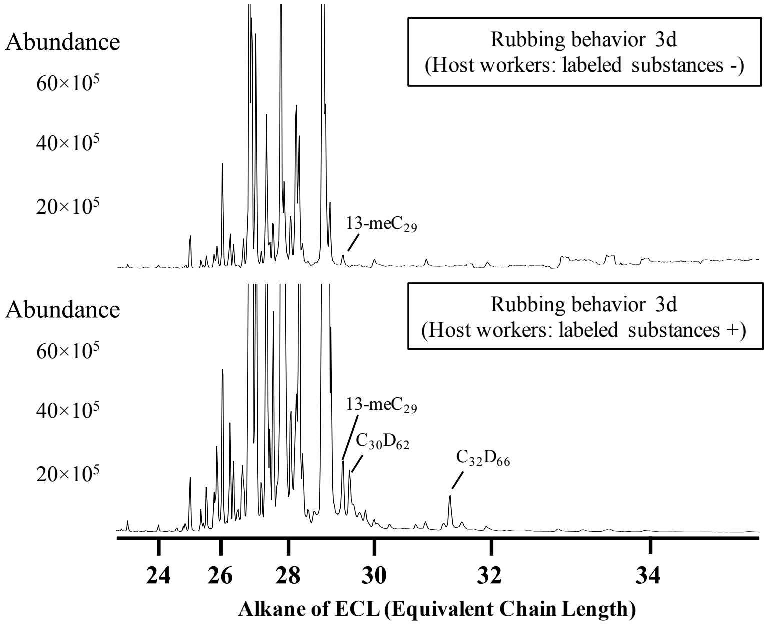


**Supplementary Figure 5. The chromatogram of labelled substances ($C_{30}D_{62}$, $C_{32}D_{66}$) from the cuticle of a newly mated *P. lamellidens* queen.** The upper panel shows the chromatograms of the newly mated *P. lamellidens* queens that exhibited rubbing behaviours towards the host workers without labelled substances, and the lower panel shows the chromatograms of newly mated *P. lamellidens* queens that performed rubbing behaviours towards the host workers with labelled substances.

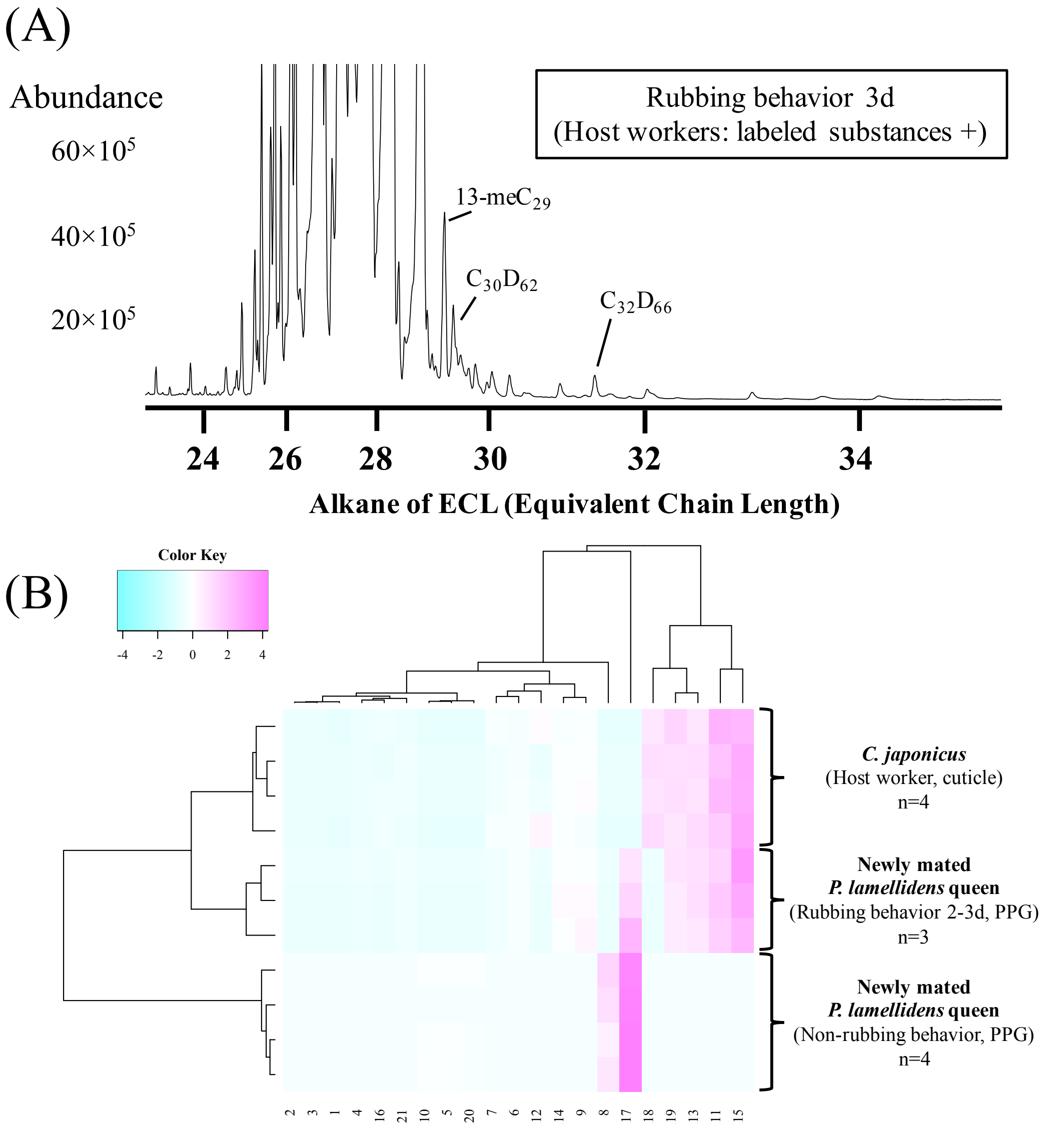


**Supplementary Figure 6. The hydrocarbons in the postpharyngeal glands (PPG) of newly mated *P. lamellidens* queens.** (A) The chromatogram of labelled substances ($C_{30}D_{62}$, $C_{32}D_{66}$) in the postpharyngeal glands of newly mated *P. lamellidens* queens. This ant exhibited rubbing behaviours towards the host workers with labelled substances. (B) Hierarchical clustering analysis of the hydrocarbons in the postpharyngeal glands and cuticle of a newly mated *P. lamellidens* queen and host workers (*C. japonicus*). This analysis was performed using the peak area value that was converted to the Z score. The number on the X-axis indicates the type of estimated hydrocarbon (see Supplementary Table 2). One of the new queens died on the second day while performing the rubbing behaviour, at which point hydrocarbons were extracted.

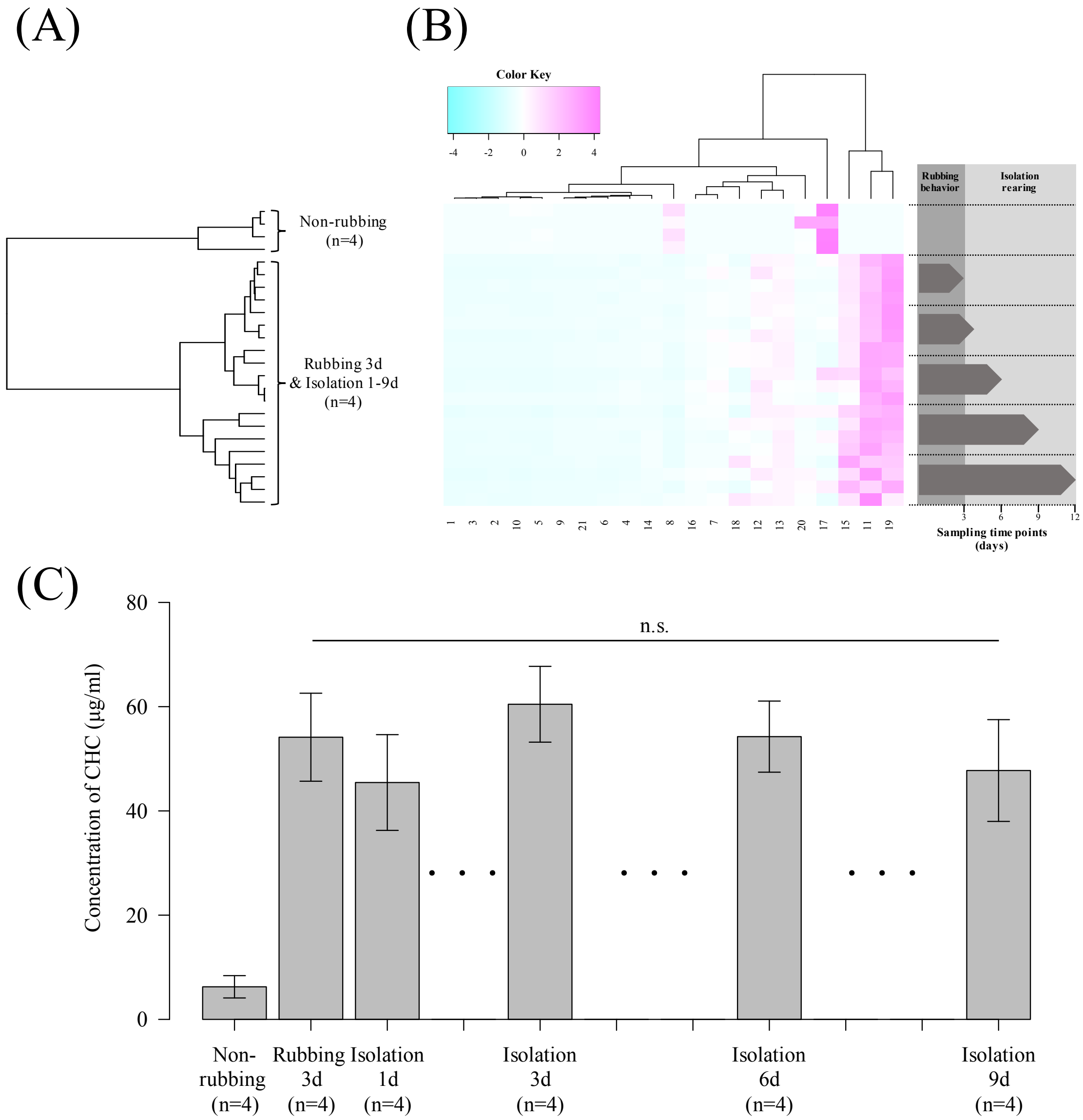


**Supplementary Figure 7. Transition of the CHC in newly mated *P. lamellidens* queens during isolation rearing.** (A) Hierarchical clustering analysis of CHC in the newly mated *P. lamellidens* queen. This analysis was performed using the peak area value that was converted to the Z score. Newly mated *P. lamellidens* queens exhibited rubbing behaviour (3d) against their hosts before being isolated from these hosts. (B) Heatmap of CHC in the newly mated *P. lamellidens* queens. This analysis was also performed using the peak area value that was converted to the Z score and used newly mated *P. lamellidens* queens that exhibited rubbing behaviour (3d) against their hosts before they were isolated from these hosts. The number on the X-axis indicates the type of estimated hydrocarbon (see Supplementary Table 2). (C) The concentration of CHC in the newly mated *P. lamellidens* queens (Mann–Whitney U test, n.s.; nonsignificant difference, n=4). The error bars indicate the standard error.

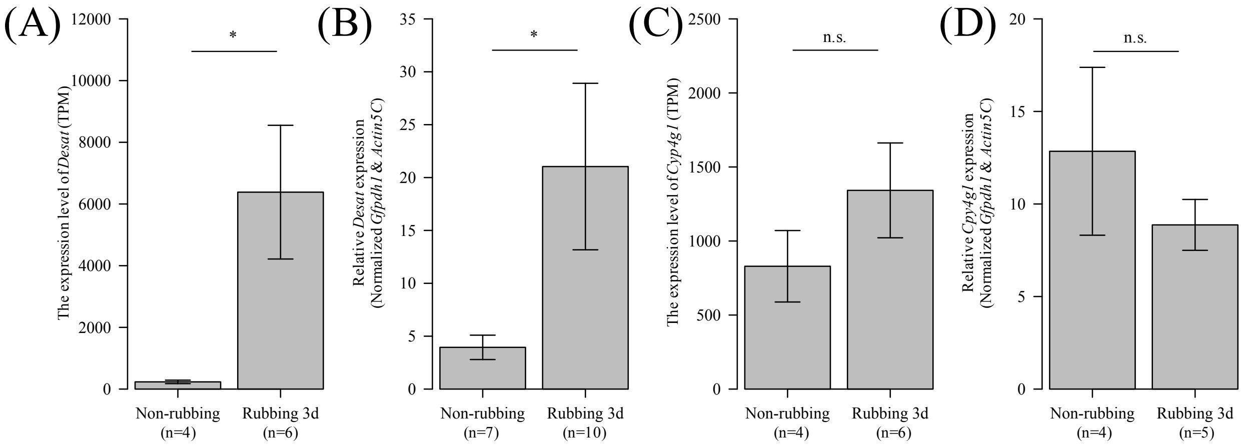


**Supplementary Figure 8. The levels of expression of CHC synthesis-related genes in fat bodies.** (A&B) The levels of desaturase (*Desat*): g34819.t1 expression in fat bodies as analysed by transcriptome (A, *; significant difference: FDR < 0.05, n=4-6) and RT–qPCR (B, *; significant difference: p value < 0.05, Welch's t test, n=7-10). (C&D) The levels of cytochrome P450 decarbonylase (*Cyp4g1*): g28547.t1 expression in fat bodies as analysed by transcriptome (C, n.s.; non-significant difference, n=4-6) and RT–qPCR (D, n.s.; non-significant difference, Welch's t test, n=4-5). All error bars indicate the standard error.

**Supplementary Movie 1. A newly mated *P. lamellidens* queen exhibits rubbing behaviour against *C. japonicus*.**
